# *In vivo* validation of novel non-invasive PHP.eB AAVs as a potential therapeutic approach for alpha-Synucleinopathies

**DOI:** 10.1101/2025.05.16.653961

**Authors:** Maria Fouka, Iraklis Tsakogias, Elena-Georgia Gialinaki, Athanasios Stavropoulos, Christiane Volbracht, Louis De Muynck, Diederik Moechars, Ronald Melki, George K. Tofaris, Leonidas Stefanis, Maria Xilouri

**Affiliations:** Biomedical Research Foundation Academy of Athens-BRFAA, Clinical-Experimental Surgery & Translational Research, Athens, Greece; Medical School, National and Kapodistrian University of Athens; 11527 Athens, Greece; H. Lundbeck A/S, Neuroscience-, Ottiliavej 9, 2500 Valby, Denmark; Janssen Research & Development, a Division of Janssen Pharmaceutica N.V., 2340 Beerse, Belgium; CEA, CNRS, Université Paris-Saclay, MIRCen, Laboratoire des Maladies Neurodégénératives, Fontenay-Aux-Roses 92260, France; Nuffield Department of Clinical Neurosciences, John Radcliffe Hospital, University of Oxford, Oxford OX3 9DU, U.K; Kavli Institute for Nanoscience Discovery, Dorothy Crowfoot Hodgkin Building, University of Oxford, Oxford OX1 3QU, U.K; School of Medicine, First Department of Neurology, National and Kapodistrian University of Athens, Greece

**Keywords:** AAV-PHP.eB, alpha-Synuclein, dopaminergic system, microRNAs, pre-formed fibrils, shRNAs

## Abstract

Parkinson’s disease (PD) is characterized by the accumulation of alpha-synuclein (aSyn) aggregates in specific brain regions, which are likely to be the disease-causing entities. Herein, we employed novel, systemically administered, brain-penetrating viral vectors (PHP.eB AAVs) in order to evaluate the potential therapeutic utility of lowering the endogenous aSyn protein burden in the aSyn pre-formed fibril (PFF)-mouse model. Such vectors expressing short hairpin RNAs or micro RNAs targeting the mouse *Snca* transcript (or respective scrambled control sequences) were intravenously administered in adult wild-type mice and two weeks later human aSyn PFFs were injected into the dorsal striatum. Following the administration of the *Snca-*targeting PHP.eB AAVs, a successful widespread viral transduction was achieved throughout the brain, accompanied by an efficient reduction of endogenous aSyn protein levels within transduced dopaminergic neurons. Intrastriatal injection of human aSyn PFFs led to the formation of pSer129-aSyn-rich cytoplasmic inclusions in brain regions connected to the PFF-injection site, nigrostriatal degeneration and relevant behavioral motor deficits, at 2.5 months post PFF-injection. Importantly, PHP.eB AAV-mediated down-regulation of endogenous aSyn reduced the accumulation of pSer129-aSyn^+^ inclusions, mitigated nigrostriatal degeneration and alleviated motor impairments. Spread of pathology to other brain regions was also attenuated.

Overall, such data highlight further the contribution of the intracellular aSyn protein load to the spread of pathology and suggest that this non-invasive delivery strategy holds promise in the research avenues for treating neurodegenerative diseases with widespread pathology, such as Synucleinopathies.

## Introduction

Parkinson’s disease (PD) is the second most prevalent neurodegenerative disorder characterized primarily by a progressive loss of dopaminergic (DA) neurons in the substantia nigra (SN) and their projections to the striatum, while other areas of the central and peripheral nervous system are also affected by the neurodegenerative process. PD is anticipated to become the fastest-growing neurological ailment, potentially affecting up to 12 million individuals by the year 2040 [1], causing grave social and economic burden to the aging societies. Current therapeutic strategies predominantly offer symptomatic relief for the motor symptoms and these benefits tend to wane after a few years. The neuronal protein α-Synuclein (aSyn) stands as the culprit of the underlying neurodegeneration, as aSyn, through its encoding gene *SNCA*, is genetically linked to familial and sporadic PD [2], while aSyn-enriched deposits are found in the remaining DA neurons either in the form of intraneuronal proteinaceous cytoplasmic inclusions named Lewy bodies (LBs) or in the form of dystrophic Lewy neurites (LNs) [3]. Consistent with the more generalized neurodegenerative process, LBs and LNs have a more widespread distribution in the nervous system [4]. Disruption in the delicate equilibrium of aSyn protein levels, including altered rates of synthesis, aggregation, release or clearance, can promote the formation and accumulation of toxic oligomeric and fibrillar species [5].

The “propagation hypothesis” posits that the spreading of aggregates is perpetuated as proteopathic seeds instruct normal aSyn to misfold and propagate from donor to recipient cells in a “prion-like” manner [6], eventually leading to neurodegeneration. The aSyn pre-formed fibril (PFF) model, founded upon the propagation hypothesis, posits that even a small quantity of preformed aSyn fibrils is sufficient to permute normal proteins into insoluble aggregates [7]. Small seeds of non-phosphorylated PFFs trigger the recruitment of endogenously expressed aSyn into pathological aggregates [8]. The focal initiation and the subsequent spread along anatomically connected structures sets the “structural template” [9]. The subsequent degeneration observed in PD-associated areas precedes the onset of motor symptoms, in line with observations in human disease [10]. This temporal characteristic enables the tracking of aggregate progression from their early formation to the point of neuronal demise. We previously demonstrated that aSyn expression level within neurons from different brain regions defines PFF seeding propensity and subsequent pSer129-aSyn accumulation [11]. Moreover, unlike aSyn overexpression-based models that result in supraphysiological levels of the protein, potentially leading to spurious neuroinflammation and neurodegeneration, total aSyn levels in the PFF model remain closer to normal physiological levels [12]. Furthermore, aSyn viral vector-based models tend to restrict the pathology temporally and spatially, falling short of the protracted phase and widespread dissemination of classical LB-like inclusion pathology [12]. Thus, the PFF model could be employed to thoroughly investigate disease initiation and progression with great temporal and spatial resolution.

Recombinant adeno-associated viral (rAAV) vectors represent an optimal platform for gene delivery, as they lack viral coding sequences and can carry therapeutic gene expression cassettes. Hence, their low immunogenicity combined with their broad range of infectivity and stable expression renders the AAVs the leading vehicle for gene therapy *in vivo* [13]. The challenge lies in delivering these vectors through non-invasive systemic routes, a critical consideration for the translation to humans on a large-scale context. However, the narrow ability of certain AAV serotypes to traverse the blood-brain barrier (BBB) and achieve adequate neuronal transduction has hindered their exploitation in the treatment of central nervous system (CNS) disorders [14]. Recently, a breakthrough has been achieved through the utilization of a Cre-recombination-based AAV targeted evolution strategy. This approach led to the identification of a novel AAV capsid variant, designated AAV PHP.eB, which differs from the AAV-PHP.B capsid by the presence of the DGT instead of the AQTLAVPFK sequence [15]. PHP.eB AAVs demonstrate enhanced neuronal tropism, and superior and more stable CNS transduction following systemic administration, attributed to their improved ability to cross the BBB, while simultaneously minimizing off-target liver transduction [15].

Herein, we employed the AAV variant PHP.eB (AAV.PHP.eB) as the selection vector for the delivery of short hairpin RNA (shRNAs) or microRNA (miRNAs) sequences targeting the endogenous mouse *Snca* gene (sh-aSyn or miR-aSyn) or respective control-scrambled sequences, prior to the intrastriatal delivery of human aSyn PFFs, to combat aSyn-related pathology. Following the intravenous administration of AAV.PHP.eB, we observed a neuronal transduction rate ranging between 15% to 40% within specific brain regions that persisted even after five months post-injection. In animals injected with sh-aSyn or miR-aSyn AAV PHP.eBs, as compared to those treated with the control-scrambled AAV PHP.eBs, we observed a noteworthy ∼ 50% reduction of endogenous aSyn protein levels within transduced nigral neurons, accompanied with decreased nigrostriatal neurodegeneration triggered by PFF injection. This amelioration of dopaminergic degeneration was reflected in the enhanced performance of the animals in the challenging beam test, indicative of an improved motor phenotype. Additionally, aggregated pSer129-aSyn levels were found decreased in both sh-aSyn+PFFs- and miR-aSyn+PFFs-treated animals in the nigrostriatal axis and in all other examined PD-related brain regions, suggesting a widespread deceleration of aSyn propagation.

Overall, such data suggest that lowering the endogenous aSyn protein load may represent an attractive approach to counteract PD-related neurodegeneration and highlight the use of AAV.PHP.eB as a promising tool for further research in non-invasive gene delivery against various brain diseases.

## Materials and Methods

### In vitro assessment of the effects of endogenous aSyn down-regulation in murine primary cortical neurons using AAV9-shRNAs or AAV1/2 microRNAs targeting the mouse Snca

Murine cortical neurons were isolated from day E14–16 embryos and cultured in Neurobasal medium supplemented with 2% B-27 supplement with antioxidants, 0.5 mM L-glutamine, 100 U/mL penicillin, and 0.1 mg/mL streptomycin (all solutions from Invitrogen, Waltham, MA, USA). Dissociated neurons were plated on 100 µg/mL poly-L-lysine coated dishes at a density of about 40,000 cells/well in 96 well plate format and transduced at days *in vitro* (DIV) 1 with two different mouse aSyn targeting AAVs. We used AAV9-Snca-shRNA (cat. # shAAV-272717) containing a 1:1 mixture of 2 mouse Snca-shRNA sequences #106636 and #272292 (#106636 5’-CCGGGTCCTCTATGTAGGTTCCAAACTCGAGTTTGGAACCTACATAGAGGACT TTTT-3’ and #272292 5’-CCGGACCAAAGAGCAAGTGACAAATCTCGAGATTTGTCACTTGCTCTTTGGTT TTTT-3’) and scramble control-shRNA (cat. # 7045) from Vector Biolabs (Malvern, PA 19355, USA) at final titers of 10×10^9, 4×10^9, and 1.6×10^9 virus particles/well and AAV1/2-Snca-3xmiRNA (cat. # GD1009-RV-M) containing three mouse Snca miRNA sequences aSyn (244): 5’-TATATTCCCAGCTCCCTCCACGTTTTGGCCACTGACTGACGTGGAGGGCTGGG AATATA-3’, aSyn (327): 5’ ATGTCTTCCAGGATTCCTTCCGTTTTGGCCACTGACTGACGGAAGGAACTGGA AGACAT-3’ and aSyn (397): 5’-TTCAGGCTCATAGTCTTGGTAGTTTTGGCCACTGACTGACTACCAAGAATGAGCCTGAA-3’ and scramble control miRNA (cat. # GD1009-RV-C) from Vigene Biosciences (Vigene Biosciences, Rockville, MD 20850, USA) at final titers of 5.5×10^7, 2.75×10^7, and 0.55×10^7 virus particles/well. 1 μM cytosine arabinoside was added to halt proliferating cells with a 50% media change at DIV4. The proportion of glia cells in the cultures was less than 10%, as assessed by an antibody against glia fibrillary acidic protein at DIV7. Transduction efficiency was estimated by counting green fluorescent protein (GFP) positive cells at DIV7: about 70-90% GFP positive cells at the two highest titers and about 40-60% GFP positive cells at the lowest titer.

At DIV7, cells were incubated with a final concentration of 1 ng/µl aSyn PFFs with a 50% media change. Another 50% media change was performed at DIV12. Cells were fixed with ice-cold 80% methanol at DIV15, labeled with pSer129-aSyn antibody (Abcam EP1536Y) and pSer129-aSyn^+^ and Hoechst fluorescence were read by Cellomics Array Scan VTI HCS Reader (ThermoFisher Scientific, Waltham, MA, USA) with a 20x objective and analyzed in ThermoScientific HCS Studio: Cellomics Scan Version 6.6.0 (ThermoFisher Scientific, Waltham, MA, USA). Seeding was measured as pSer129-aSyn Spot Total Intensity/viable cell count. Methanol fixation impaired the GFP signal, which analysis was omitted at DIV15. Viable cell count was performed at a separate re-scan only counting Hoechst positive cell bodies from cells that were alive before fixation, based on nuclear size and fluorescent intensity. Each condition was measured in 5 technical replicates (wells) per culture experiment, with each well imaged in 16 fields. Cells from parallel plates transduced with AAVs at the indicated titers were harvested at DIV 15 to determine knockdown efficiency and measuring mouse aSyn levels by ELISA from LSBio (cat. # LS-F6284, LSBio, Shirley, MA 01464, USA) following manufacturer’s instructions and total protein concentration using the bicinchoninic acid (BCA) Protein Assay Kit from Pierce as per manufacturer’s instructions (ThermoFisher Scientific, Waltham, MA, USA). Each condition was measured in 3 technical replicates (wells) per experiment.

### Animals

Eight-week-old male wild-type (WT) C57Bl/6 mice (20–28g body weight) were housed at the Animal House Facility of the Biomedical Research Foundation of the Academy of Athens (Athens, Greece) in individually ventilated cages under specific pathogen–free conditions, on a 12-hour light/12-hour dark cycle and ad libitum access to food and water. All experimental procedures performed were approved by the Ethical Committee for Use of Laboratory Animals of the Biomedical Research Foundation Academy of Athens (60898 and 478434 license numbers) and in accordance with the ARRIVE guidelines and the EU Directive 2010/63/EU for animal experiments.

### Production of AAVs

All AAV vectors utilized in the study were generated by SIRION Biotech (Table 1). These AAV plasmids expressed nuclear-localized GFP (NLS-GFP) driven either from the constitutive CAG promoter (AAV-CAG-eGFP) together with an artificial short hairpin RNA (two different sh-aSyn sequences, #272292 and #106636) or the constitutive CBA promoter (AAV-CBA-eGFP) bearing an artificial micro-RNA (miR-aSynx3) targeting the mouse *Snca* to down-regulate endogenous aSyn expression or control vectors with respective scrambled shRNA (scramble shRNA, sh-SCR) or micro-RNA (miNegx3, miR-SCR). AAV vectors were packaged with the PHP.eB capsid that can cross the BBB [15]. The sh-aSyn-272292-treated group displayed a marked decrease in mobility and a rapid decline in body weight, indicative of severe toxicity. Consequently, these animals were ethically euthanized ten days following the injection and all analyses presented thereof originated using the sh-aSyn-106636 sequence.

**Table 1.**
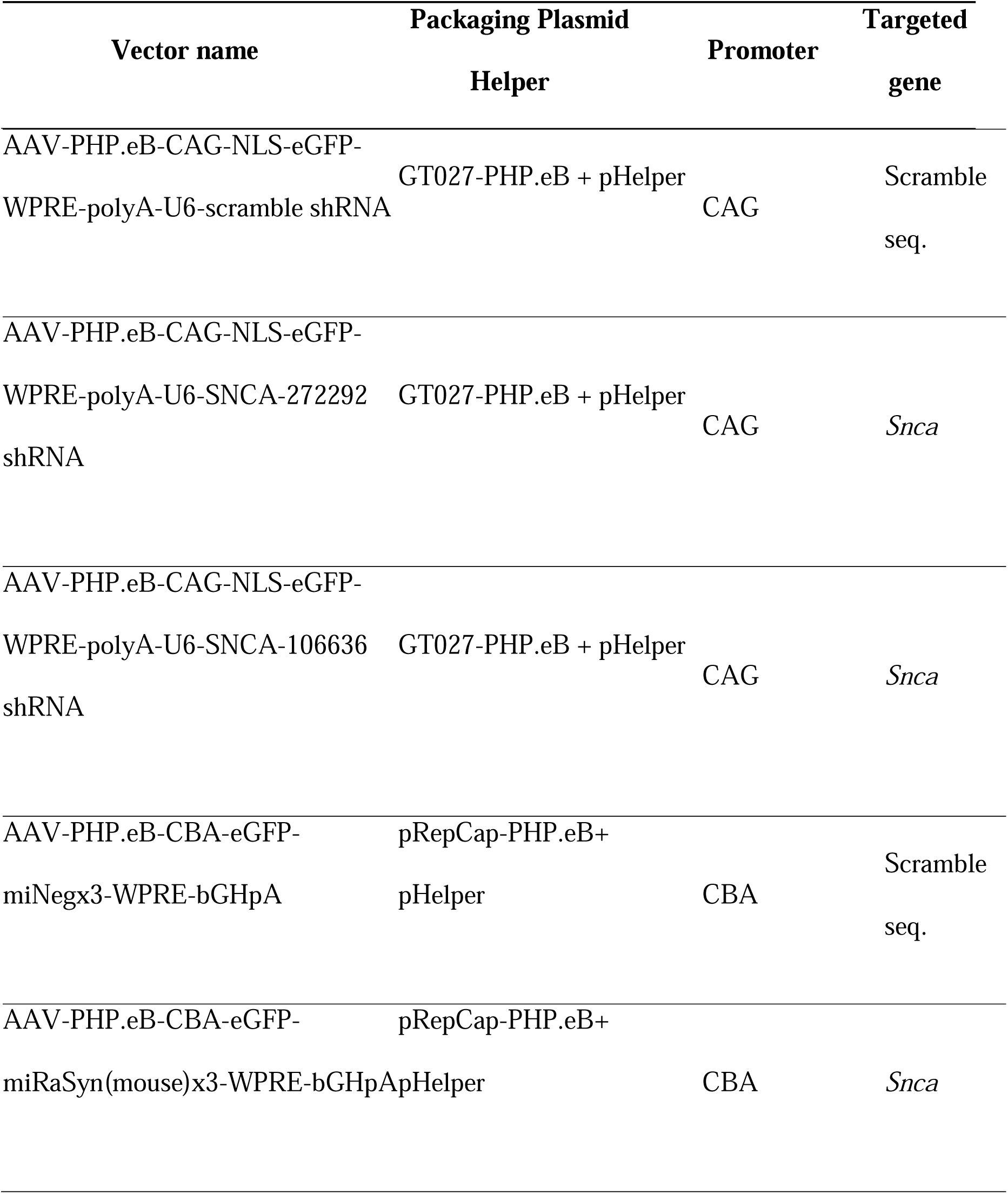
Table of the AAV-PHP.eB vectors’ characteristics employed in intravenous administration.

### Intravenous (i.v.) Tail Vein Injection

Intravenous administration of AAV PHP.eB vectors was performed by injection into the tail vein of twelve-week-old mice. Mice were placed in a heated recovery chamber (40°C ± 2) for 10 min prior to each injection to dilate veins. After, the conscious mice were placed in a restrainer, and then a 29 gauge needle was inserted into a lateral tail vein as previously described [16], [17]. Mice were then injected with 200 μL of AAV-PHP.eB (2E^11^ viral genome (vg)) or PBS as control for 30 to 40 s. The same person performed all injections in the study in one day.

### Preparation of human aSyn pre-formed fibrils (PFFs)

Human wild-type aSyn pre-formed fibrils (haSyn PFFs) were generated in accordance with the established protocol as described previously [18]. Briefly, human wild-type aSyn was expressed in E. coli BL21 DE3 CodonPlus cells and purified as described before [19]. The obtained monomeric aSyn was endotoxin free (Pierce LAL Chromogenic Endotoxin Quantification Kit). Fibril assembly was initiated by incubating monomeric aSyn (200µM) in 50 mM Tris–HCl, pH 7.5, and 150 mM KCl at 37°C with continuous agitation in an Eppendorf Thermomixer set at 600 rpm for a duration of 7 days [20]. The assembly reaction was monitored by thioflavin T binding employing a Cary Eclipse Fluorescence Spectrophotometer (Varian Medical Systems Inc.) set to excitation and emission wavelengths of 440 nm and 480 nm, respectively. The morphology of the resulting fibrillar assemblies and characteristic proteolytic profile were assessed through transmission electron microscopy following negative staining with a 1% uranyl acetate solution and SDS-PAGE analysis and Coomassie staining [20]. The generated fibrils were subjected to two rounds of centrifugation at 15,000 g for 10 minutes each and were subsequently re-suspended in PBS. The concentration of PFFs was adjusted to 350 μM in PBS. These fibrils were further fragmented to an average length of 40–50 nm by subjecting them to sonication for 20 minutes in 2 mL eppendorf tubes, utilizing a Vial Tweeter powered by an ultrasonic processor UIS250 v (250 W, 2.4 kHz; Hielscher Ultrasonic) [21]. The fragmented fibrils were aliquoted (6 μL) in 0.5 mL Eppendorf tubes, flash frozen in liquid nitrogen, and subsequently stored at −80°C until their intended use.

### Surgical procedures

All surgical procedures were performed under isoflurane (Abbott, B506) anesthesia. Animals were given 5 mg/kg carpofen for analgesia. After placing the animal into a stereotaxic frame (Kopf Instruments, USA), 5 μg of human artificial PFFs were injected unilaterally into the right striatum using the following stereotaxic coordinates relative to bregma: + 0.5 mm anteroposterior, −1.8 mm mediolateral, and two depths dorsoventral −3.2 and −3.4 mm from the skull, according to the mouse stereotaxic atlas [22]. Injections were performed using a finely pulled glass capillary (diameter of approximately 60–80 μm) attached to a Hamilton syringe with a 22-gauge needle at the rate of 0.1 μl/15s. Between target depths, a 5-min interval was followed before being slowly withdrawn from the brain. The unilateral PFF injections were performed two weeks following the systemic delivery of the PHP.eB AAVs (Suppl. Fig. 1A).

### Behavioral analysis

To assess motor performance and coordination, the *challenging beam* test was performed 3 months after i.v. injection, in accordance with a previously published protocol [23]. Briefly, mice were placed on a beam consisting of four 25-cm long frames of decreasing width (3.5, 2.5, 1.5 and 0.5 respectively), placed onto two inverted home cages positioned at the narrower extremity of the beam. A detachable metallic grid is placed atop the beam. Mice were trained to transverse the beam from the widest to the narrowest frame over a two-day period, with the grid omitted during the trial phase. The trials concluded when the mouse inserts one of its forelimbs into the home cage, which was located at the end of the beam to encourage them to cross it in the desired direction. On the third day, the grid was placed on top of the beam and the animals were videotaped throughout five trials, which were then viewed by a blind observer to detect any recorded errors or slips through the grid. Errors included limb slips through the grid during forward motion. Motor impairment is correlated with an increased number of paw slips through the mesh grid in forward motion. The time to cross the beam, the total number of steps and the ratio of errors per step were also counted. Total slips were averaged across the five trials for each animal.

The *rotarod assay* was employed to assess motor coordination and balance. Mice are placed on a horizontally oriented, rotating cylinder, which is low enough not to injure the animal, but high enough to induce avoidance of fall. Rodents naturally try to stay on the rotarod and avoid falling to the ground. This assay is performed three times on the same day, with each trial being 2 hours apart. Mice are evaluated on the length of time they can stay on the rotarod.

### Tissue preparation

Mice were anesthetized with isoflurane and perfused intracardially with phosphate-buffered saline (PBS; Thermo Fisher Scientific, 70011036), followed by ice-cold fixative 4% PFA (Sigma Aldrich, P6148) under a constant flow rate using a pump. The brains were post-fixed overnight at 4°C in the same preparation of paraformaldehyde and then transferred to 15% sucrose in PBS overnight, before being incubated in 30% sucrose until freezing. For snap freezing, brains were immersed into frozen 2-methylbutane (Isopentane) at −45°C for 30-50 s and stored at −80°C. Free floating sections of 30 μm increments, encompassing the whole rostro-caudal axis of the brain were collected, using a Leica cryostat at −25°C.

### Immunohistochemistry

General immunohistochemical stainings were carried out in free-floating sections as described [24]. Briefly, tissue sections were washed 3 times in PBS (3×10 min). The selected sections were incubated for 10 min in a 10% MeOH/3% H_2_O_2_ solution (in PBS), followed by blocking of the non-specific sites with 5% normal goat serum (NGS) in PBS. The primary antibodies against mouse tyrosine hydroxylase (TH) at a dilution of 1:5000 for the striatal and 1:3000 for the nigral sections and against pSer129-aSyn at a dilution of 1:10.000 for striatum- and substantia nigra-containing sections and 1:7500 for the rest brain regions, were solubilized in 2% NGS (in PBS) and sections were incubated with the respective antibodies for 24 h. The secondary biotinylated rabbit antibody was subsequently used (1:3000; 60 min in NGS 2% in PBS), followed by incubation with the ABC solution (Elite ABC kit, Vector) and staining with 3’-diaminobenzidine (DAB, Vector Laboratories), according to manufacturer’s instructions. The sections were afterwards mounted at Superfrost^+^ slides (VWR) and dehydrated.

### Immunofluorescence

Tissue sections were washed 3 times in PBS (3×10 min), and then permeabilized and blocked for 1 hour in blocking solution (5% NGS, 0.1% Triton-X in PBS) at 25°C. Sections were incubated with primary antibodies (2% NGS, 0.1% Triton-X in PBS) shown in Table 2, overnight at 4°C. Following three washing steps of 10 min each with PBS, sections were incubated with the secondary antibodies and DAPI for 1h at 25 °C and washed again as above. Finally, the sections were rinsed with PBS and mounted on poly-D-lysine coated glass slides.

**Table 2.**
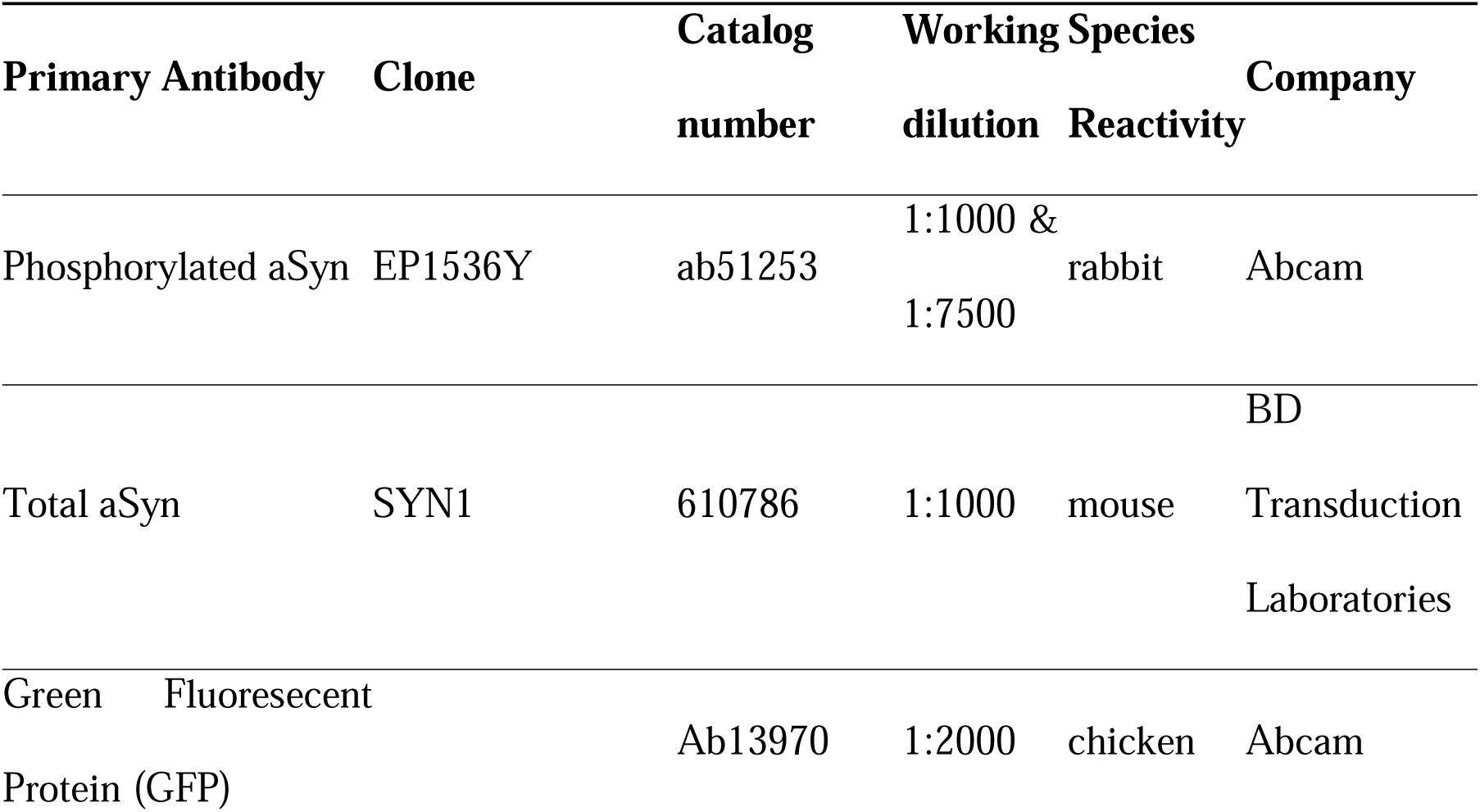
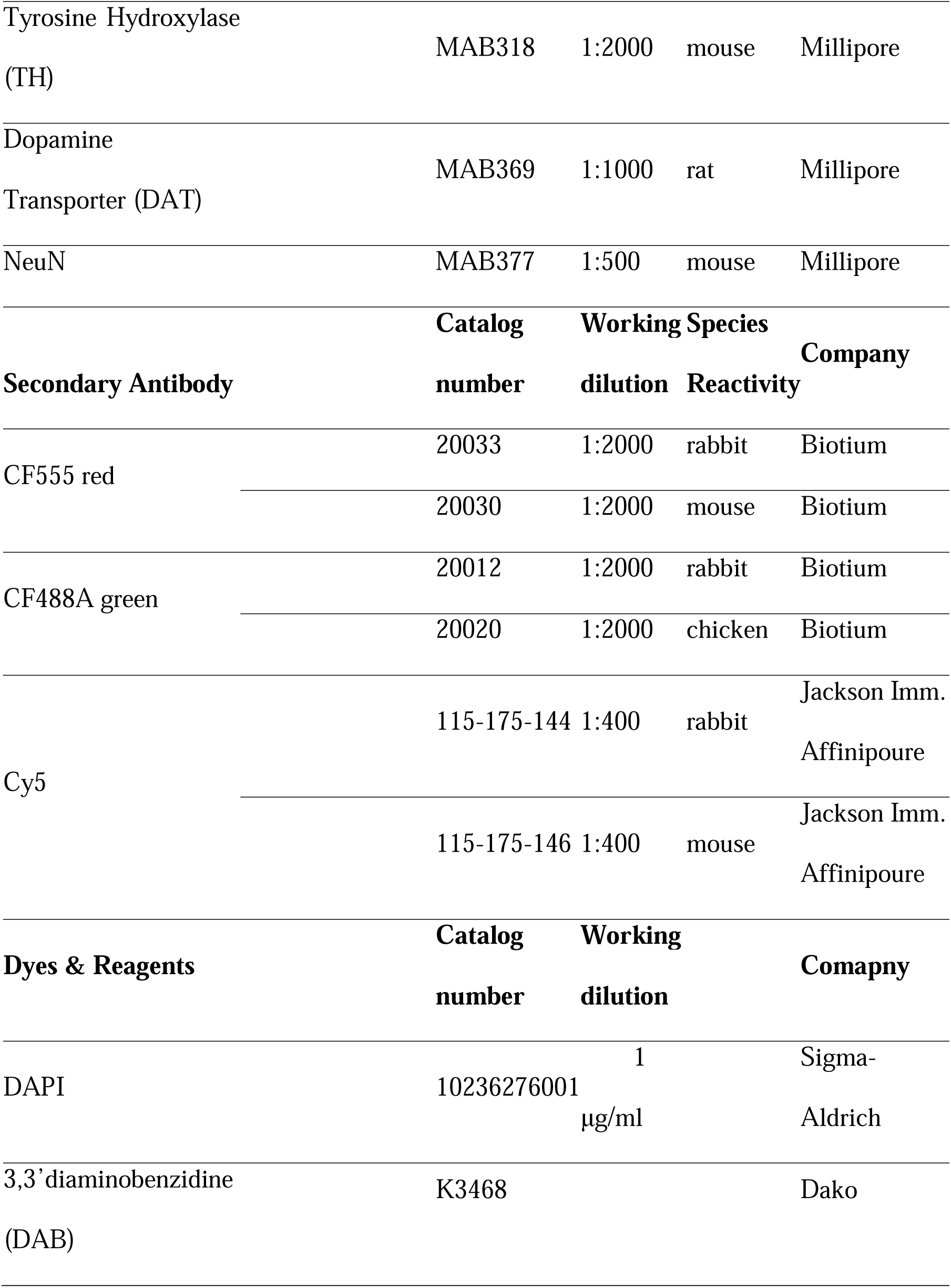
List of antibodies used for immunofluorescence (IF) and immunohistochemistry (DAB) stainings.

### Image analysis and quantification

The ImageJ and Imaris (ver 10.0) software were used for image analysis. For the mapping of the spreading of aSyn pathology, coronal sections spanning from the olfactory bulb to the substantia nigra in interslice intervals of approximately 180 μm, and 120 μm for substantia nigra, were stained for pSer129-aSyn and developed using DAB. Immunostained tissue was visualized on a Leica DMRA2 light microscope and captured with a QImaging top-mounted camera. The Allen Brain Atlas (Allen Institute, USA; http://atlas.brain-map.org/) was used as a reference for assigning neurons to regions. For the quantification of pSer129-asyn positive area, color segmentation plugin on ImageJ was used, allowing us to pixel cluster the pSer129-aSyn positive signal in a semi-automated way.

For the immunofluorescence intensity measurement of dopamine transporter (DAT) and the area of pSer129-aSyn, images were split into distinct color channels and converted to binary after a threshold was applied to eliminate the background auto-fluorescence. For analysis of average fluorescence intensities for assays such as ascertaining the intensity of aSyn signal in the cells of the substantia nigra, the image analysis program Imaris was used, utilizing the “surfaces” tool. Imaris was also utilized to calculate transduction efficacy of the AAVs using the “spots” function, by creating spots for all GFP and NeuN (neuronal marker) or TH (dopaminergic neuron marker) and then using “colocalize spots” to find out how many cells had been penetrated by the GFP-expressing virus.

### Stereology

Tyrosine hydroxylase (TH)-positive nigral neuron density was estimated by an unbiased stereological quantification method utilizing the optical fractionator principle embedded within the Stereo Investigator v10.0 software (MicroBrightField Bioscience, Magdeburg, Germany), a motorized stage (MAC5000, Ludl Electronic Products, Ltd., NY, USA) and a top-mounted camera (QImaging, MicroBrightField Bioscience, Magdeburg, Germany) with the light microscope (Leica, DMRA2, Wetzlar, Germany). Every fourth section throughout the whole rostro-caudal axis of the substantia nigra was immunostained for TH using the 3,3’-diaminobenzidine (DAB). Τhe region of interest was outlined with a Leica 2.5×0.07 objective (50601) and counting was performed using a Leica 63×1.3 glycerol immersion objective (506192). A random start and systematic sampling was applied. Counting parameters were adjusted to achieve at least 100 counts per hemisphere. A single blinded investigator performed all quantifications and analysis. A coefficient of error (Gundersen, m=1) of 0.1 was accepted.

### Densitometry of TH^+^ dopaminergic striatal terminals

Striatal tyrosine hydroxylase-positive fiber density was measured by densitometry using every sixth coronal section of the animal’s striatal tissue, covering the whole rostro-caudal axis of the striatum. The images were converted into 8-bit binary; the regions of interest were outlined, and the density of the TH-positive terminals was measured. The measured values were corrected for non-specific background staining by subtracting values obtained from the cortex. The data are presented as means from all animals and expressed as the ratio to the contralateral side.

### Statistical analysis

GraphPad Prism 8 software was used for all statistical analysis. Data were tested for normality using the Shapiro–Wilk test. For comparisons between two groups, either a two-tailed Student’s *t*-test or Mann–Whitney test was applied, depending on the data distribution. For comparisons involving multiple groups with two independent variables, a two-way ANOVA was conducted, followed by Sidak’s or Tukey’s multiple comparison tests for post-hoc analysis. Statistical significance was set as \**p* < 0.05, \*\**p* < 0.01, \*\*\**p* < 0.001, \*\*\*\**p* < 0.0001 and data are presented in graphs as mean ± SEM. At least n = 3-4 animals/group for the shRNA-PHP.eB and n = 5-7 animals/group for the miRNA-PHP.eB AAVs were included in the analyses presented. Raw data are available upon reasonable request.

## Results

### Prevention of seeded aSyn aggregation by AAV-mediated knockdown of endogenous mouse *Snca* in murine primary neuronal cultures

To assess the impact of endogenous aSyn down-regulation οn the phosphorylation levels and seeding of endogenous rodent aSyn, we initially validated two different AAV-based knockdown approaches targeting the endogenous *Snca* transcript in mouse primary cortical cultures, using the human aSyn PFF model. We used an AAV9 construct containing two different mouse aSyn shRNA sequences #272292 and #106636 (AAV9-Snca-shRNA from Vector Biolabs, cat. #shAAV-272717) and one AAV1/2 construct containing three artificial microRNA sequences targeting mouse *Snca* (AAV1/2-Snca-3xmiRNA from Charles River, catalog number: GD1009-RV-M). Both AAVs contained eGFP as reporter. One-day old (DIV1) primary cortical cultures were transduced with the AAV9-Scna-shRNA or scramble control shRNAs using final titers of 1×10^10, 4×10^9, and 1.6×10^9 virus particles/well and analyzed for endogenous mouse aSyn levels by ELISA at 15 days *in vitro* (DIV15). We observed sufficient knockdown of 100%, 98%, and 77%, respectively using AAV9-Scna-shRNA (Fig. 1A, left panel). Similar treatment of primary cortical cultures transduced with the AAV1/2-Snca-miRNA constructs using final titers of 5.5×10^7, 2.75×10^7, and 5.5×10^6 virus particles/well revealed a knockdown in mouse *Snca* expression of 99%, 90% and 51%, respectively (Fig. 1A, right panel).

**Figure 1.**
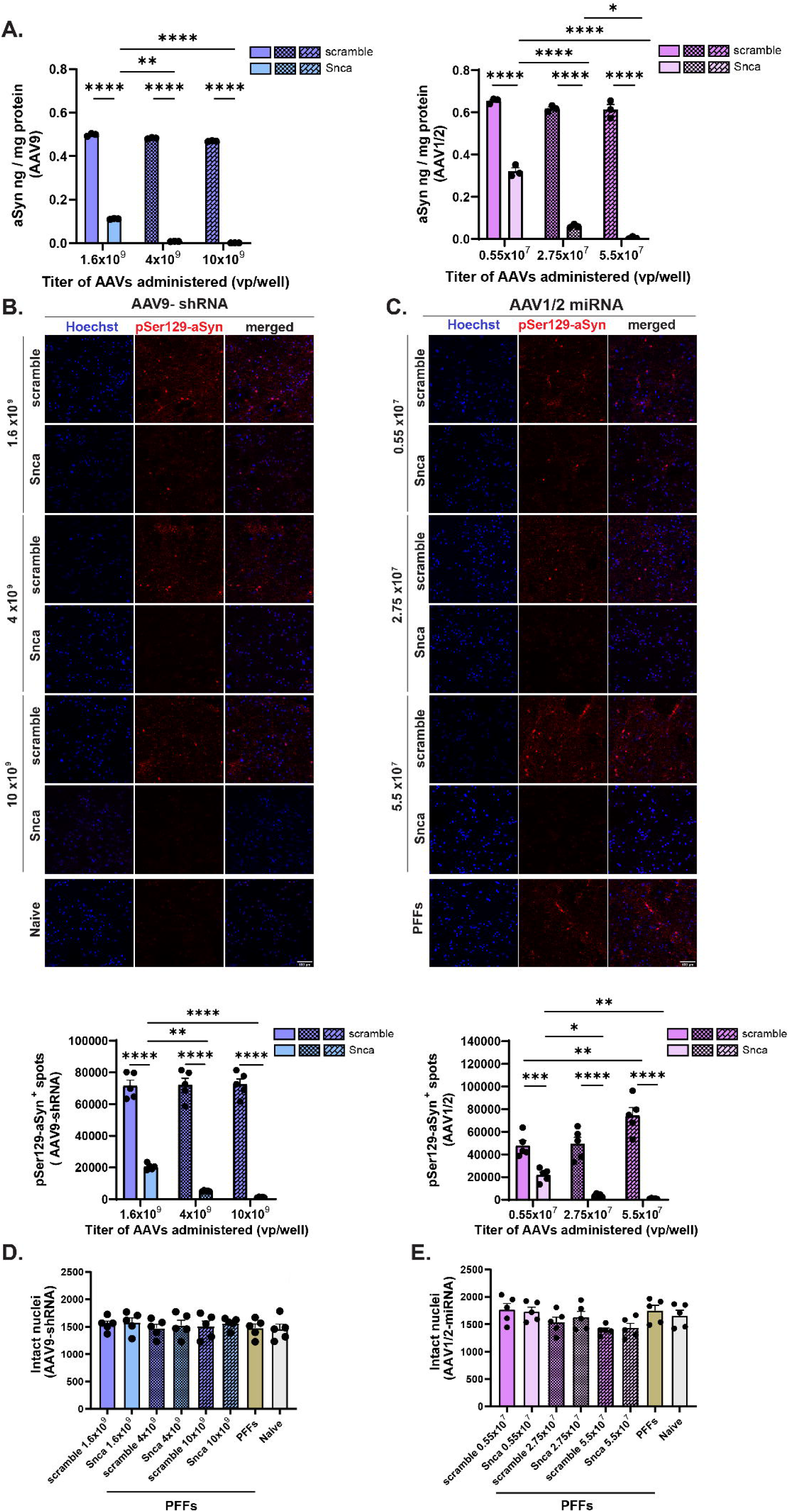
AAV mediated aSyn knockdown prevented seeded aSyn aggregation *in vitro*. Mouse primary cortical neurons were transduced at DIV 1 with *Snca* knockdown AAV9-shRNA or AAV1/2-miRNA and respective scramble control sequences in the titers indicated. (**A**) At DIV 15, significant reduction in aSyn levels in AAV9-shRNA-(*left*) or AAV1/2-miRNA-(*right*) treated cultures was detected, as compared to the respective SCR-control-treated cultures. Alpha-synuclein protein levels were detected by ELISA and data are presented as mean ± SD of three (3) independent experiments. Comparisons were made using Two-way ANOVA followed by Tukey’s post hoc test, asterisks indicate significance (*p < 0.05, ** p < 0.01, ****p < 0.0001). (**B-C**) At DIV 7, neurons were treated with haSyn PFFs (1 ng/µl) and processed for pSer129-aSyn immunocytochemistry to reveal seeded aSyn aggregation at DIV 15. Representative confocal images (20x objective, *upper panels*) of nuclei stained with Hoechst (in gray), and pSer129 alpha-synuclein immunoreactivity (in red) and respective graphs (*bottom panels*) of AAV9-shRNA-(*left*) or AAV1/2-miRNA-(*right*) treated cultures are shown. Scale bar = 580 µm. Quantifications of pSer129-aSyn immunoreactivity are shown as Spot Total Intensity/viable cell count as mean ± SD of five (5) independent experiments. Comparisons were made using Two-way ANOVA with Tukey’s post hoc test, asterisks indicate significance (*p < 0.05, ** p < 0.01, *** p < 0.001, ****p < 0.0001). (**D**) Quantifications of the number of intact nuclei in AAV9-shRNA-(*left*) or AAV1/2-miRNA-(*right*) treated cultures, in the presence of haSyn PFFs at DIV 15. A one-way ANOVA followed by Tukey’s post hoc test was used to assess differences in the number of healthy nuclei among groups with different titers within the same AAV serotype. No significant differences were observed in the number of healthy nuclei between any groups in both AAV1/2 and AAV9 constructs (n=5).

Subsequently, we tested the impact of endogenous mouse aSyn down-regulation on pSe129-aSyn levels by treating cells with human aSyn PFFs (1 ng/µl) at DIV7. The effects of *Snca* knockdown on seeded aSyn aggregation were analyzed by quantification of spots positive for pSer129-aSyn at DIV15. A reduced seeded aSyn aggregation by 98%, 93% and 71%, was observed in cultures treated with 1×10^10, 4×10^9, and 1.6×10^9 virus particles/well AAV9-Scna-shRNA, respectively (Fig. 1B). Likewise, highly reduced seeded aSyn aggregation was observed in cultures treated with the two highest titers of AAV1/2-Snca-3xmiRNA by 99% and 92%, respectively (Fig. 1C). In accordance with 51% knockdown of mouse aSyn, we observed 54% reduction of seeded aSyn aggregation with the lowest titer of AAV1/2-Snca-3xmiRNA. No toxicity was observed at the applied titers of any AAV construct (in the presence of PFFs), as assessed by measuring the numbers of intact nuclei in all conditions (Fig. 1D).

### Intrastriatal injection of haSyn PFFs increases aSyn phosphorylation at Ser129 within the injected striatum and impairs nigrostriatal innervation

To establish aSyn-related pathology in the murine brain, human aSyn PFFs were unilaterally inoculated into the right dorsal striatum of wild-type mice, a PD-vulnerable region that is also receptive to the uptake of aSyn PFFs [25]. Our primary objective was to validate our in-house PFF model’s capacity to impact PD-vulnerable brain regions and, in doing so, to demonstrate a high level of face validity by mirroring characteristics observed in human PD.

We first assessed the levels of pSer129-aSyn-positive immunofluorescent signal within the striatum, the area of PFF injection (Fig. 2A-B), given that this residue is recognized as the predominant pathological modification of aSyn commonly associated with Lewy bodies and Lewy neurites [26]. At the two and a half months’ time interval, pSer129-aSyn^+^ inclusions were evident along the rostrocaudal axis of the ipsilateral striatum and, to a lesser extent, in the contralateral hemisphere (Fig. 2A upper row & B). We next analyzed the impact of PFF-injection on the striatal dopaminergic fiber integrity by assessing the dopamine transporter (DAT) intensity, a reliable marker of presynaptic dopaminergic terminal loss. The PFF-injected hemisphere displayed a modest decrease in DAT mean fluorescent intensity (MFI) when compared both with the non-injected hemisphere and the respective PBS-injected hemisphere of the control group (Fig. 2C). It is noteworthy that the observed attenuation in DAT intensity was predominantly localized in close proximity to the injection site, which coincided with the pronounced presence of PFFs. This spatial correlation suggests that the loss of the DAT levels is intricately associated with increased aSyn protein burden, which is further supported by a negative correlation coefficient between pSer129-aSyn^+^ area and the DAT MFI of the PFF-injected hemisphere relative to its respective contralateral hemisphere (Fig. 2D).

**Figure 2:**
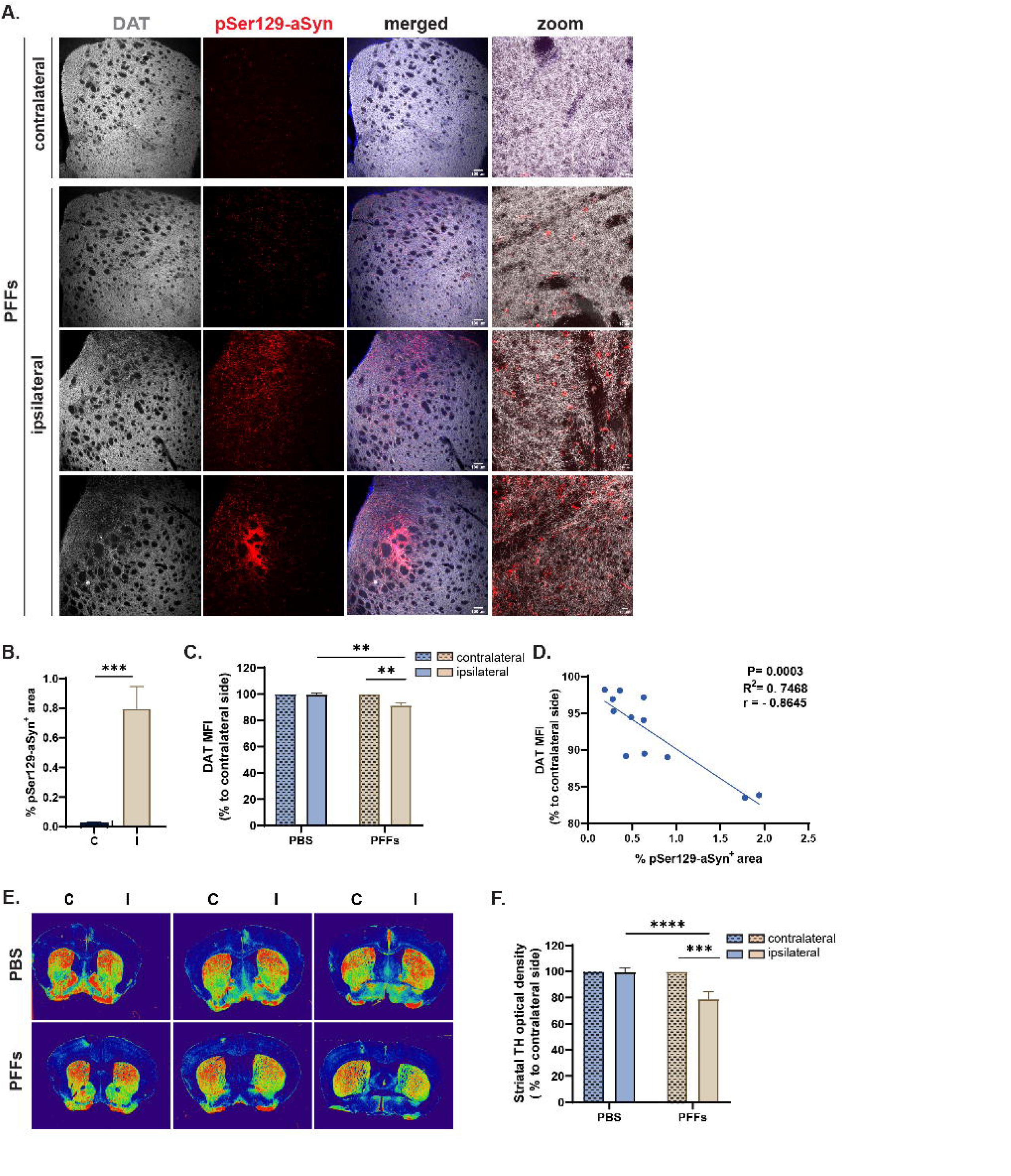
Intrastriatal injection of haSyn-PFFs increases Ser129-aSyn phosphorylation within the injected striatum and impairs nigrostriatal innervation. (**A**) Representative immunofluorescence images of striatal sections with antibodies against dopamine transporter (DAT) and pSer129-aSyn in PFF-treated animals, at 2.5 months post injection. Pathological pSer129-aSyn^+^ inclusions are present throughout the rostra-caudal axis of the PFF-injected striatum (ipsilateral, three different levels are shown at rows 2-4), whereas scarce pSer129-aSyn^+^ signal can also be detected in the contralateral side; however, to a lesser extent. The right panel displays a higher magnification of DAT and intracellular pSer129-aSyn^+^. (**B-C**) Quantification of the percentage of pSer129-aSyn^+^ area (B) and of the mean fluorescence intensity (MFI) of DAT (C) in the striatum of PFF-treated animals, at 2.5 months post injection. Data are presented as the mean percentage ± SEM relative to the corresponding contralateral side. (**D**) Negative linear correlation (r=-0.8645) between the pSer129-aSyn % area and the mean DAT intensity (***, p=0.0003, R^2^=0.7468, Pearson correlation coefficient). (**E-F**). DAB immunostaining in coronal striatal sections of PFF-treated animals using tyrosine hydroxylase (TH) antibody (E) and quantification of the relative TH^+^ optical density expressed as the percentage of the non-injected contralateral side (F). Error bars represent mean ± SEM, with n=4-6 animals/group. Comparisons were made using unpaired t-tests, asterisks indicate significance (**p < 0.01, ***p < 0.001, ****p < 0.0001). Scale bar: 100 µm and 10 µm (zoom).

To further corroborate the dopaminergic striatal pathology, we assessed the optical density of tyrosine hydroxylase (TH) positive dopaminergic terminals in DAB-stained coronal striatal slices (Fig. 2E). A 21% decrease in dopaminergic terminals was observed in the PFF-injected hemisphere, as compared to the non-injected contralateral counterpart (Fig. 2F). Notably, degeneration was predominantly present at the dorsal part of the striatum, corresponding to the site of PFF injection. This observation probably implies that the pathological degeneration does not extend throughout the striatum within the relatively short timeframe of 2.5 months post-injection.

### Intrastriatal injection of haSyn PFFs evokes Ser129-aSyn phosphorylation within host mouse nigral neurons and impairs dopaminergic neuron survival and motor performance

Increased pSer129-aSyn^+^ signal was also detected in the ipsilateral substantia nigra (SN) implicating the retrograde transfer of haSyn PFFs from the striatum to the SN and the templating of the endogenous protein at 2.5 months post injection (Fig. 3A-B). No pSer129-aSyn positive signal was detected in both hemispheres of the PBS-injected cohort (data not shown). To ascertain whether the reduction in the striatal dopamine terminals depicted in Fig. 2 correlated with concurrent dopaminergic cell loss, we conducted an unbiased stereological quantification of dopaminergic neurons within the substantia nigra pars compacta (SNpc, Fig. 3C-D). The fibrils elicited an 18% reduction in the number of TH^+^ neurons within the SNpc of the ipsilateral hemisphere (Fig. 3D), consistent with the corresponding decline in TH immunoreactivity in the nigrostriatal terminals (Fig. 2E-F). In PBS-treated animals, TH striatal intensity appeared to be preserved and the numbers of nigral dopaminergic neurons closely aligns with the reported numbers for the C57BL/6 mouse strain [27]. This observation indicates that the surgical and stereotactic procedures employed for the injection of exogenous haSyn PFFs did not result in noticeable tissue and/or cellular damage that could significantly enhance cellular uptake, thus creating an artificial enhancement of susceptibility.

**Figure 3.**
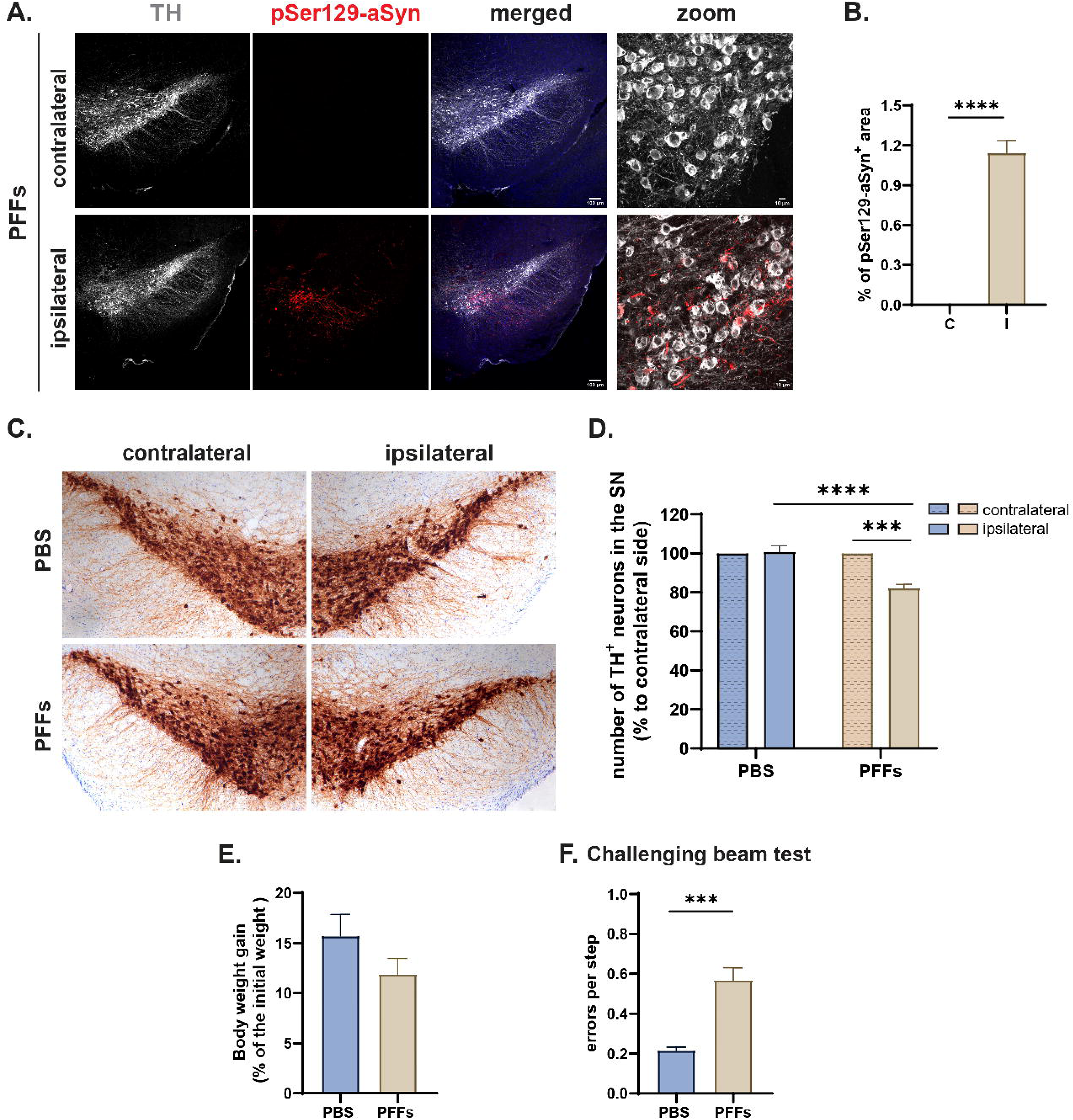
HaSyn-PFFs intrastriatal delivery evokes Ser129-aSyn phosphorylation within dopaminergic neurons, impairs cell survival and motor performance. (**A**) Representative immunofluorescence images of midbrain sections with antibodies against TH and pSer129-aSyn in PFF-treated animals at 2.5 months post injection depicting the presence of pSer129-aSyn^+^ aggregates in the ipsilateral substantia nigra (SN). Intracellular pSer129-aSyn^+^ inclusions within nigral dopaminergic neurons are shown in higher magnification in the right panel. Scale bar: 100 µm, 10 µm. (**B**) Quantification of the percentage of pSer129-aSyn^+^ area in the SN of PFF-treated animals, at 2.5 months post injection. (**C-D**) DAB immunostaining in coronal midbrain sections of PBS- and PFF-treated animals using TH antibody (C) and stereological quantification of TH^+^ neurons in the injected and non-injected hemisphere of PBS- and PFF-treated animals (D), at 2.5 months post injection with n=3-6 animals/group, Error bars represent mean ± SEM. Comparisons were made using two-way ANOVA, Šidák correction, asterisks indicate significance, **p < 0.01, ***p < 0.001, ****p < 0.0001. (**E**) Body weight gain of PBS- and PFF-treated animals during the course of study (**F**) Quantification of errors per step measured by the challenging beam test (***p < 0.001; n = 6 animals/group, t-test). Error bars represent mean ± SEM.

In order to have a rudimentary gauge of the potential surgical and PFF inoculation-evoked toxicity, animal body weight was recorded at weekly intervals. Although the PFF-injected group collectively displayed a trend towards attenuated weight gain compared to PBS-injected animals, this trend did not reach statistical significance (Fig. 3E). To evaluate the capacity of haSyn PFF inoculation to faithfully reproduce a parkinsonian-like phenotype within the designated timeframe, we scrutinized parameters pertaining to motor behavior, motor coordination, and balance. Challenging beam test has been previously proved successful at detecting motor disturbances of the hind limbs [23]. PFF-administration was indeed efficacious in inducing a motor impairment phenotype at 2.5 months post-injection, as evidenced by a greater frequency of grid slips during the challenging beam traversal in the PFF-injected group compared to the control PBS-injected mice (Fig. 3F).

### Widespread transduction of murine CNS neurons following intravenous administration of shRNA-or miRNA-GFP-PHP.eBs, at three months post-systemic delivery

In order to investigate the utility of the systemic administration of AAV-PHP.eB viral vectors as successful gene delivery tools into the CNS, we systemically administered, through the lateral tail vein, 200 μL of AAV-PHP.eB-GFP viral vectors containing 2E^11^ viral genome (vg) copies/animal into 8-week-old male C57/Bl6 mice. The animals were administered the sh-aSyn (CAG promoter) or the miR-aSyn (CBA promoter) PHP.eB constructs that were shown to efficiently down-regulate the endogenous murine *Snca* in primary cortical cultures or respective SCR controls (Fig. 1A). The animals were euthanized at a 3-month interval subsequent to the intravenous administration. This specific temporal juncture was selected to facilitate the adequate expression of the transgenes introduced and to evaluate the enduring effects of the therapeutic intervention. The GFP and shRNA or microRNA aSyn-targeting sequences lie downstream of a constitutive CAG or CBA promoter, respectively, which has been proposed to ensure sustained, long-term expression across a spectrum of cell types [28] [29]. The systemic administration of the viral vectors, coupled with the utilization of the versatile and robust promoters, has led to the transduction of a noticeable proportion of neurons (marked with the specific marker NeuN) throughout the entire brain (Fig. 4A, C, F).

**Figure 4:**
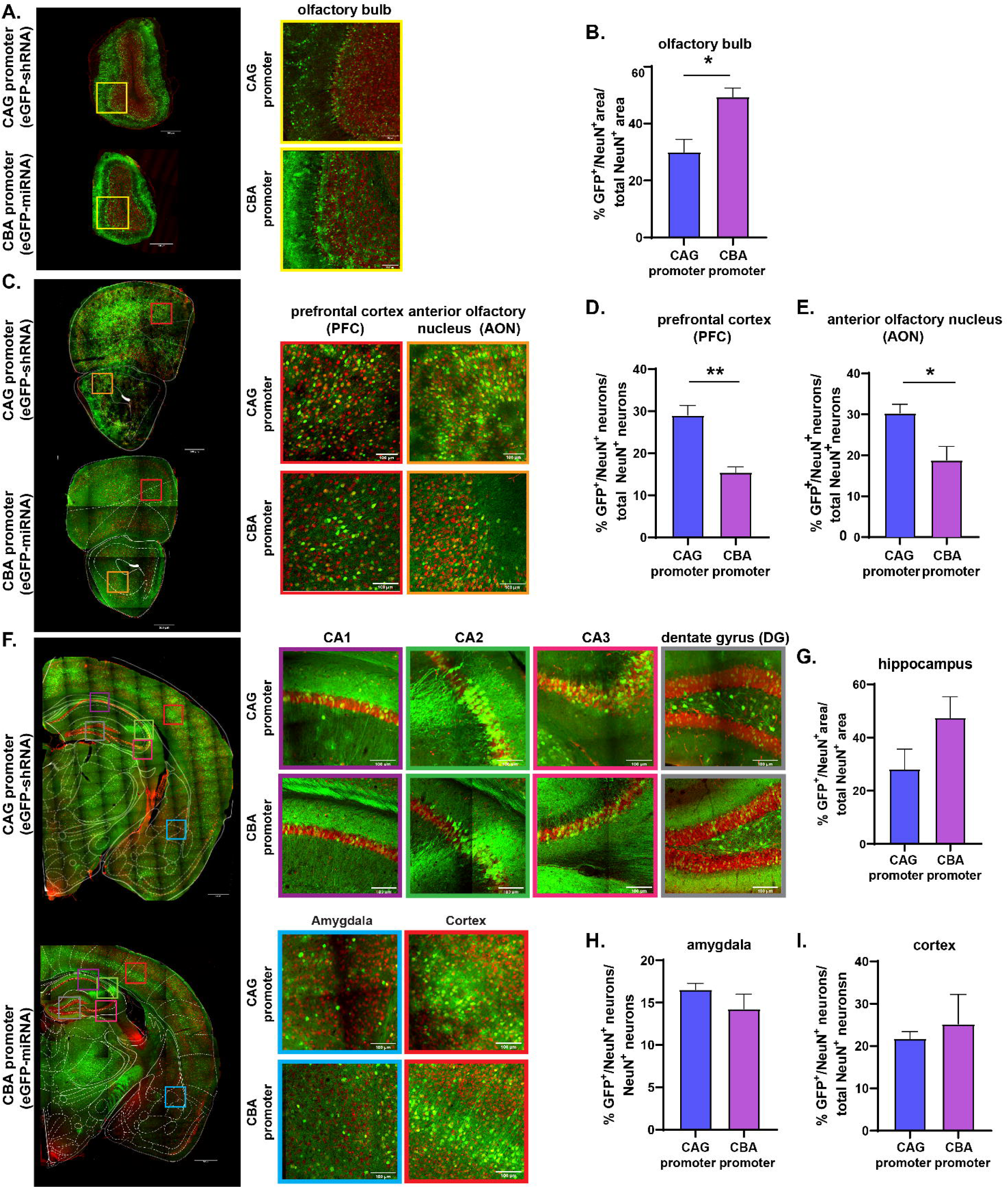
Widespread transduction of murine CNS neurons three months following intravenous administration of GFP-PHP.eBs. **(A, C, F)** Representative tile scans obtained through confocal microscopy and superimposed with mouse brain atlas grids, depicting GFP expression at different rostro-caudal coordinates of shRNA-GFP (CAG promoter, *upper panels*) and miRNA-GFP (CBA promoter, *bottom panels*) PHP.eB viral vectors. The contralateral to haSyn PFF injection sides were utilized for the image acquisition and quantitative analysis shown herein. Scale bar: 500 µm. Enlarged images, marked by colored squares, highlight specific brain regions: olfactory bulb (*black*), prefrontal cortex (PFC, *red*), anterior olfactory nucleus (AON, *orange*), hippocampal regions CA1 (*yellow*), CA2 (*green*), CA3 (*pink*), dentate gyrus (DG, *grey*), and amygdala with cortex (*blue*). Scale bar: 100 µm. (**B, D-E, G, H-I**) Quantification of the transduction efficacy of AAV-PHP.eBs expressing GFP under the control of CAG or CBA promoter assessed by quantifying the percentage of GFP^+^ area or GFP^+^ cells co-localizing with NeuN^+^ cells for both promoters. Results are presented as the percentage of GFP^+^ area relative to total NeuN^+^ area (**B, G**), or the percentage of GFP^+^/NeuN^+^ co-localized cells relative to the total number of NeuN^+^ neurons (**D-E, H-I**). (*p < 0.05; **p < 0.01, n = 3-4 animals/group, t-test). Error bars represent mean ± SEM.

In alignment with recently published findings, we observed robust transduction in the glomerular layer of the olfactory bulb, ranging between 30-50% in the PHP.eBs harboring the CBA and CAG promoter, respectively (Fig. 4B). Nevertheless, precise quantification of the transduced neurons within the olfactory bulb (Fig. 4B) and the hippocampus (Fig. 4G) proved challenging due to the densely packed arrangement of NeuN^+^ cells. To overcome this, we computed the % ratio of both GFP^+^ and NeuN^+^ area against the total NeuN^+^ area. Moving to the prefrontal cortex (PFC) and the anterior olfactory nucleus (AON), the transduction efficacy reached ∼ 30% and 15-20% in each of these regions, for the PHP.eBs harboring the CAG and CBA promoter, respectively (Fig. 4D-E). A notable GFP transduction in the hippocampus, particularly in the CA2 sub-region, was observed with both AAVs, with the PHP.eB harboring the CBA promoter exhibiting the highest percentage, reaching ∼ 45% (Fig. 4F-G). The data obtained by the co-localized GFP^+^/NeuN^+^ versus the total NeuN^+^ area quantifications in PFC and AON brain areas (Fig. 4C-E) versus the hippocampus (Fig. 4F-G), probably indicate a region-specific tropism of the two different promoters. In the amygdala (Fig. 4H) and associated cortex (Fig. 4I) cells exhibiting dual labeling for GFP and NeuN accounted for ∼ 15% of the total neuronal population (Fig. 4H-I). Collectively, both CAG and GBA promoters utilized in the study are ubiquitous promoters that drive high levels of gene expression, as previously reported [30].

Notably, only the contralateral to the haSyn PFF-injection sides were utilized for the image acquisition and quantitative analysis shown in Fig. 4 to circumvent any potential toxic effects evoked by the PFF-injection. Importantly, the numbers of TH^+^ dopaminergic neurons in the contralateral sides of the PBS/PFF group (Suppl. Fig. 1B) were similar to the numbers in the respective sides of the AAV+PFF-injected groups, indicating the non-toxic profile of the viruses utilized in the mouse dopaminergic system (Suppl. Fig. 1C-D). Finally, in order to have a rudimentary gauge of a potential toxicity evoked by the systemic delivery of the PHP.eBs, and/or the stereotactic delivery of the PFFs, animal body weight was recorded at weekly intervals. An increase in body weight gain was detected in shaSyn-PHP.eB-PFF-treated animals versus the untreated (WT) counterparts, while no significant alterations were apparent in the miRNA-PHP.eB-injected animals (Suppl. Fig. 1D).

### AAV-PHP.eB viral vectors expressing shRNAs or microRNAs against mouse *Snca* effectively target substantia nigral dopaminergic neurons and mitigate total aSyn protein levels

To comprehensively evaluate whether the aforementioned AAV-PHP.eB viral vectors could selectively target nigral dopaminergic neurons, we performed immunofluorescent staining, coupled with image analysis and quantification, using antibodies targeting TH (marker for dopaminergic neurons) and GFP, as a marker for the respective AAV-PHP.eB viral vectors. Our analysis revealed strong co-localization of TH^+^ nigral neurons with GFP marker throughout the rostal-caudal axis (ten sections per animal, with a 4-section interval) of the substantia nigra (Fig. 5A). Quantification of TH^+^/GFP^+^ co-localized neurons revealed that, at three months post administration, approximately 30% (CAG promoter) or 44% (CBA promoter) respectively, of dopaminergic neurons demonstrated stable GFP expression (Fig. 5B).

**Figure 5.**
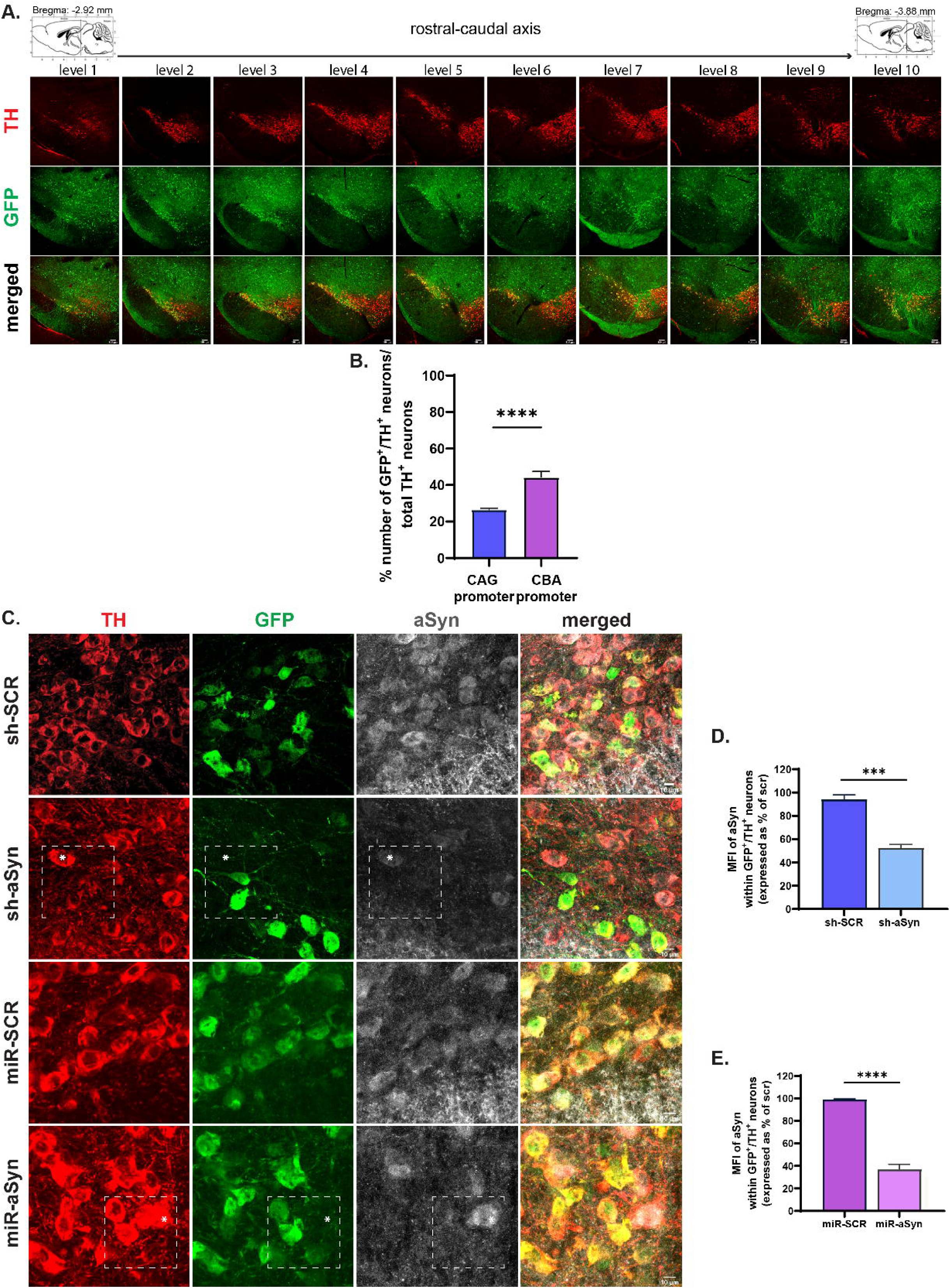
AAV-PHP.eBs bearing mouse *Snca*-targeting shRNAs or microRNAs transduce dopaminergic neurons and reduce aSyn protein levels. **(A)** Representative immunofluorescence images of midbrain sections stained with antibodies against TH and GFP, illustrating the transduction efficiency of AAV-PHP.eB-miRNAs in substantia nigra along the rostral-caudal axis, three months post intravenous administration. Scale bar: 100 μm. **(B)** Quantification of transduction efficacy of AAV-PHP.eBs expressing GFP under the control of CAG or CBA promoter in the substantia nigra measured by the percentage of GFP^+^ cells that colocalize with TH^+^ cells. n = 4-8 animals/AAV-PHP.eB subtype. **(C)** Representative immunofluorescence images of midbrain sections with antibodies against TH, GFP and aSyn illustrating aSyn levels, within GFP^+^ dopaminergic neurons transduced with AAV-PHP.eB co-expressing GFP with either shRNAs (upper panel) or microRNAs (lower panel) targeting endogenous mouse *Snca*. Asterisks indicate non-transduced GFP^-^/TH^+^ neurons that manifest increased aSyn signal. Scale bar: 10 µm. **(D-E)** Quantification of mean fluorescence intensity (MFI) of aSyn within transduced dopaminergic neurons with AAV-PHP.eB co-expressing GFP with shRNAs (D) or microRNAs (E) expressed as a percentage relative to scr injected animals. (***p < 0.001; ****p < 0.0001, n = 3-4 animals/group, t-test). Error bars represent mean ± SEM.

To conduct a thorough assessment of the potential capacity of these AAVs to effectively down-regulate endogenous aSyn protein expression, we conducted immunofluorescence analyses with a focus on the transduced (GFP^+^) dopaminergic (TH^+^) neurons, alongside with measurements of aSyn protein levels (Fig. 5C). Notably, sh-aSyn AAV or miR-aSyn AAV expression elicited a 48% (Fig. 5D) or 63% (Fig. 5E) reduction in endogenous aSyn protein expression levels, as compared to the respective controls sh-SCR- or miR-SCR-transduced TH^+^-GFP^+^ cells. Thus, the sh-aSyn or miR-aSyn AAVs successfully reduce nigral aSyn expression, thereby indicating the utility of these AAVs as a tool to ascertain the contribution of endogenous aSyn to the pathogenesis of PD and related Synucleinopathies.

### AAV-PHP.eB-mediated down-regulation of endogenous mouse aSyn ameliorates pSer129-aSyn pathology and dopaminergic neurodegeneration evoked by haSyn-PFF inoculation

To evaluate prospective therapeutic capabilities, we elected to integrate these AAVs into our established haSyn PFF model, which faithfully recapitulates a parkinsonian-like phenotype, as illustrated in Fig. 2 and 3. Considering these data, our investigation aimed to ascertain whether such decline in endogenous murine aSyn protein levels could effectively impede the protracted course of haSyn PFF-evoked neurodegeneration. To obtain a first glimpse on the effect of the successful down-regulation we conducted immunofluorescence (IF) staining targeting pSer129-aSyn and computed the percentage of the area exhibiting positive staining (Fig. 6A-F). The attenuation of endogenous aSyn levels yielded a concomitant reduction in the extent of the pSer129-aSyn^+^ area, encompassing both the pars compacta (Fig. 6C & E) and pars reticulata (Fig. 6D & F) subdivisions of the substantia nigra.

**Figure 6.**
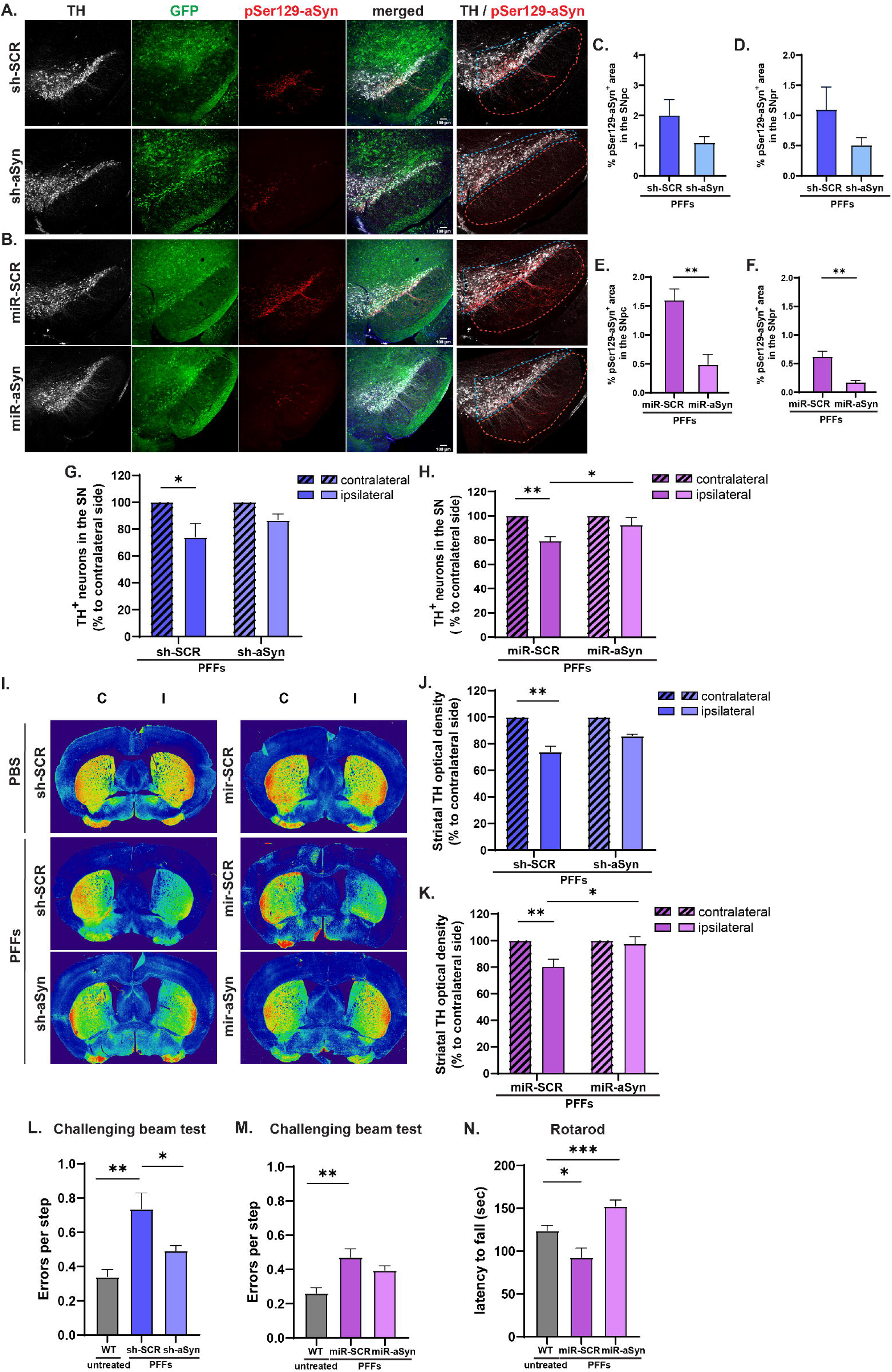
AAV-PHP.eB-mediated down-regulation of endogenous mouse aSyn ameliorates pSer129-aSyn pathology and dopaminergic neurodegeneration evoked by haSyn-PFFs. **(A-B)** Representative immunofluorescence images of midbrain sections stained with antibodies against TH, GFP and pSer129-aSyn illustrating pSer129-aSyn^+^ signal within the mouse nigral dopaminergic neurons, at 2.5 months post-PFF inoculation in animals systemically administered with sh-aSyn or sh-SCR (A) and miR-aSyn or miR-SCR (B) AAV-PHP.eBs. Higher magnification images verifying the presence of intracellular pSer129-aSyn^+^ inclusions within SNpc (substantia nigra pars compacta, blue dotted line) and SNpr (substantia nigra pars reticulate, orange dotted line) TH^+^ neurons are shown in the right panels. Scale bar: 100 μm. (**C-F**) Quantification of the percentage of pSer129-aSyn^+^ area in the SNpc (C, E) and SNpr (D, F) of animals injected with sh-aSyn or sh-SCR (C-D) and miR-aSyn or miR-SCR (E-F) AAV-PHP.eBs, at 2.5 months post-PFF administration. (**p < 0.01, n = 3-4 animals/group, t-test). Error bars represent mean ± SEM. **(G-H)** Stereological quantification of TH^+^ nigral neurons in the PFF-injected (*ipsilateral*) and non-injected (*contralateral*) hemisphere, assessed at 2.5 months post-PFF inoculation in animals systemically administered with sh-aSyn or sh-SCR (G) and miR-aSyn or miR-SCR (H) AAV-PHP.eBs. **(I-K)** TH-DAB immunostaining in coronal striatal sections (I) and quantifications of the relative TH^+^ optical density of animals injected (I, *ipsilateral*) and non-injected (C, *contralateral*) hemisphere, assessed at 2.5 months post-PFF inoculation in animals systemically administered with sh-aSyn or sh-SCR (J) and miR-aSyn or miR-SCR (K) AAV-PHP.eBs. All data shown in G, H, I, K are expressed as the percentage of the non-injected contralateral side (**p < 0.01, n = 4-5 animals/group, t-test). Error bars represent mean ± SEM. **(L-M)** Quantification of errors per step measured by the challenging beam test in all groups (***p < 0.001; n = 6 animals/group, t-test). Error bars represent mean ± SEM. **(N)** Mice were tested at increasing speeds in a rotarod apparatus and the number of cumulative falls was recorded. Quantification of mean value obtained for each animal in all three trials (**p < 0.01; n = 7-11 animals/group, One-way ANOVA followed by Tukey’s post hoc test).

To further corroborate the ameliorating effect of the aSyn down-regulation and correlate it with the preservation of the dopaminergic substantia nigra neuron integrity, we performed a stereological quantification of nigral TH^+^ neurons. As depicted in Fig. 6G & H, endogenous mouse aSyn down-regulation by either sh-aSyn or miR-aSyn AAV-PHP.eBs yielded a mild, yet discernible, mitigation of the extent of neuronal depletion induced by PFFs, which reached statistical significance in the miR-aSyn treated animals, as compared to miR-SCR-injected ones (Fig. 6H). To further assess the impact of these pathological aggregates on the integrity of the nigrostriatal axis we utilized densitometric evaluation of the neuronal terminals in the injected and the non-injected striatum of all animals, at 2.5 moths post PFF injection (Fig. 6I-K). Densitometric analysis of striatal dopaminergic fiber density by TH immunoreactivity revealed that similar to what was observed in the nigra, sh-aSyn or miR-aSyn AAV-PHP.eB-treated animals exhibited a less pronounced reduction in dopaminergic terminals evoked by PFF treatment, as compared to the respective controls (Fig. 6I-K), with the effect of miR-aSyn displaying a higher statistical power (Fig. 6K).

Collectively, endogenous aSyn silencing ameliorated the effects on dopaminergic system integrity evoked by haSyn PFF inoculation and impeded the propagation of aSyn-related pathology within the nigrostriatal pathway. To establish a correlation between these therapeutic effects and the observed motor dysfunction triggered by haSyn PFF injection (Fig. 3F), mice were subjected to the challenging beam test. PFF injection in mice treated with the control scrambled AAVs (sh-SCR or miR-SCR) evoked an increased frequency of errors during their traversal of the beam, whereas endogenous aSyn silencing (sh-aSyn or miR-aSyn treated animals) ameliorated this motor deficit (Fig 6L-M). Further supportive of this conclusion are the data obtained from the rotarod test where the miR-aSyn-PFF group performed significantly better than the miR-SCR-PFF group (Fig. 6N). Interestingly, assessment of GFP expression driven by the CAG promoter within the nigrosriatal pathway at 5 months post i.v. delivery, revealed a sustained GFP expression at comparable levels to those at 2.5 months, underscoring the stability achieved by the PHP.eB vectors and thereby bolstering their potential as a long term gene delivery platform (Suppl. Fig. 2A-B). Furthermore, a reduction in pSer129-aSyn^+^ area was evident in both striatum (Suppl. Fig. 2C-D) and nigra (Suppl. Fig. 2E-F) of mice treated with sh-aSyn-PFF as compared to the sh-SCR-PFF animals (Suppl. Fig. 2C-F). Additionally, evaluation of the nigrostriatal terminal integrity (Suppl. Fig. 2G-H) and related motor behavior (Suppl. Fig. 2I) elicited by PFF inoculation, uncovered a partial rescue in the sh-aSyn-PFF animals, compared to sh-SCR-PFF ones. However, more animals need to be incorporated in the study to draw reliable conclusions.

### Systemic administration of AAV PHP.eB harboring shRNAs or miRNAs against the endogenous murine *Snca* mitigates pSer129-aSyn-related pathology at interconnected brain regions, upon intrastriatal haSyn PFF-inoculation

Following unilateral haSyn PFF injection, a gradient of aSyn-related pathology was evident that spanned areas beyond the nigrostriatal axis, at 2.5 months post-PFF administration, presumably facilitated through trans-synaptic anatomical connections. We next assessed whether endogenous mouse aSyn down-regulation through the systemic administration of the aSyn-targeting AAVs could mitigate the progressive spread of pathological aSyn species along anatomically connected brain structures. Thus, we focussed on brain areas displaying direct synaptic connections to the nigrostriatal pathway such as the prefrontal cortex, hippocampus, cortex and amygdala, in which aSyn pathology appears at different stages of the disease as defined by the Braak stages [4]. Importantly, all these regions exhibited positive immunoreactivity for pSer129-aSyn as detected 2.5 months post-PFF injection (Fig. 7). The relative transmission pattern and regional abundance of aSyn^+^ inclusions in the different PHP.eB AAV-injected groups were depicted in the form of heat maps, where a regression of pSer129-aSyn-related pathology relative to scrambled controls, is evident in all areas analyzed in both sh-aSyn and miR-aSyn groups (Fig. 7A). In particular, decreased pSer129-aSyn^+^ area was detected in both contralateral (control) and ipsilateral (PFF-inoculated) sides of the sh-aSyn and miR-aSyn prefrontal cortex that reached statistical significance only in the miR-aSyn ipsilateral hemisphere (Fig. 7B-C). In the hippocampus (Fig. 7D-E) and cortex (Fig. 7F-G), a significant decline of pSer129-aSyn^+^ area was evident in both sides of sh-aSyn and miR-aSyn treated animals. Lastly, in the amygdala, a region characterized by extensive connections with various CNS nuclei with a particular emphasis on the dorsomedial striatum, a significant decrease was detected in both sides of miR-aSyn treated animals (Fig. 7 H-I). The effect was less prominent and may be attributed to the smaller sample size used in the sh-aSyn group (Fig. 7 H-I). Both the thalamus and hypothalamus exhibited minimal pSer129 aSyn^+^ signal (Fig. 7 J-K) at the ipsilateral PFF-injected hemispheres in all groups and were not analyzed due to suboptimal detection with the color segmentation tool.

**Figure 7.**
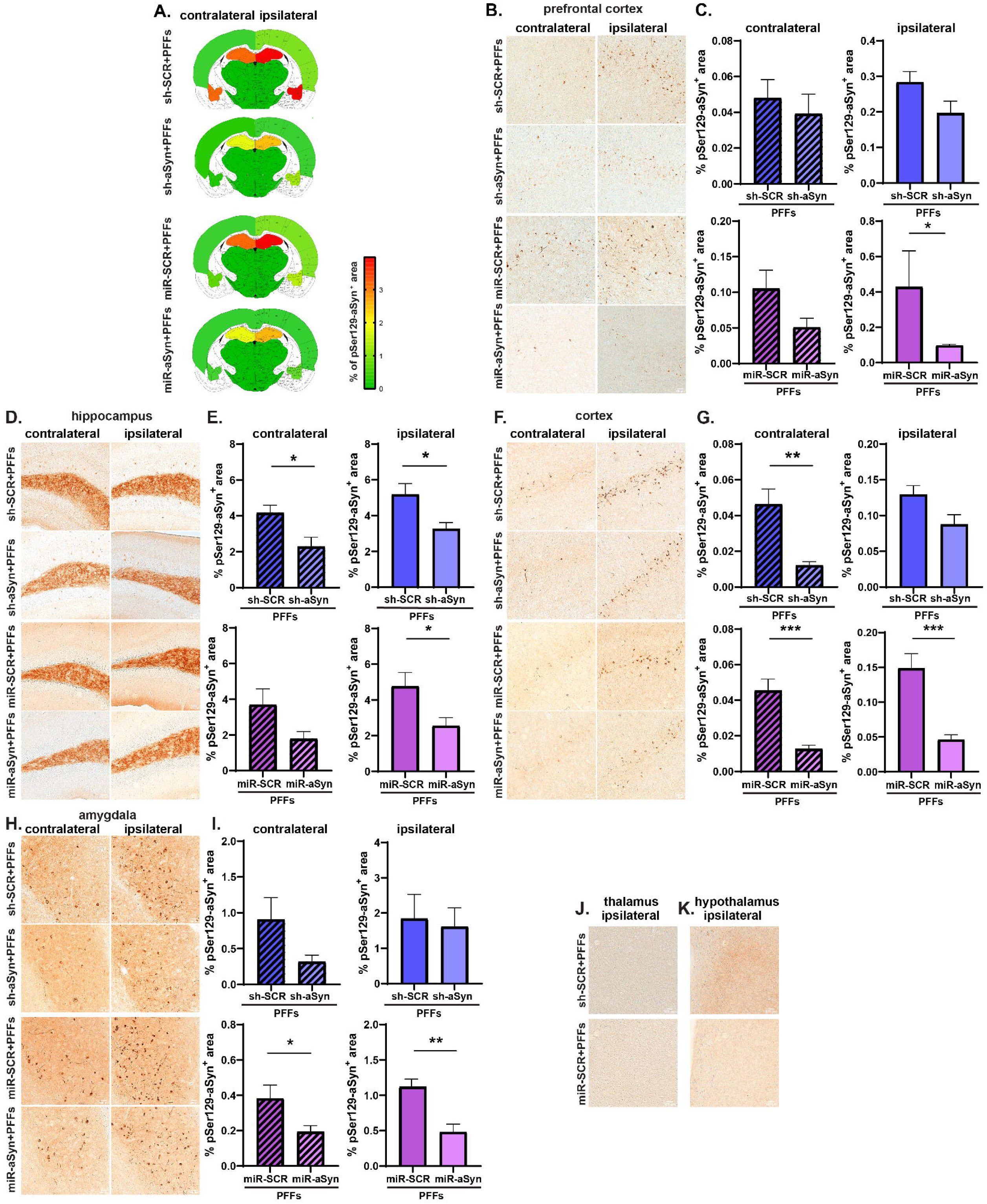
Systemic administration of murine *Snca*-targeting AAV-PHP.eBs mitigates pSer129-aSyn-related pathology at interconnected areas, upon intrastriatal PFF-inoculation. **(A)** Heat maps of aSyn-related pathology at contralateral/ipsilateral sides of shRNA or miRNA AAV-PHP.eB-injected animals, 2.5 months following unilateral PFF inoculation. Semi-quantitative analysis of the percentage of the pSer129-aSyn^+^ area graded from negative (green) to most abundant (red, 6) and color-coded onto heat maps. **(B-I)** Representative DAB-stained sections against pSer129-aSyn and the relative quantifications of pSer129-aSyn^+^ area at pre-frontal cortex (PFC) (B-C), hippocampus (D-E), cortex (F-G) and amygdala (H-I). Reduced pSer129-aSyn^+^ area is evident in both hemispheres of the sh-aSyn or miR-aSyn-treated groups, although to a different extent. In all statistical comparisons unpaired t-test was performed, *p < 0.05; **p < 0.01; *** p < 0.001, n = 4-6 animals/group. **J**) Representative DAB-stained sections against pSer129-aSyn at the ipsilateral thalamus and hypothalamus demonstrating lack of aSyn-related pathology in all groups at 2.5 months post-PFF injection.

## Discussion

Herein, we aimed to investigate the effects of limiting the endogenous aSyn expression in a well-established mouse model of PD induced by the intracerebral injection of human aSyn PFFs. The novelty of our approach lies in the strategic utilization of the novel AAV.PHP.eB viral vectors featuring a CAG or a CBA promoter. This combination facilitated the intravenous administration of the viral vector, while also enabling its widespread distribution throughout the entire brain. This method allowed us to specifically target and down-regulate in wide areas of the brain the expression of murine aSyn, a pivotal player in the pathogenesis PD and related synucleinopathies. By capitalizing on the non-invasive AAV.PHP.eB systems, we achieved a comprehensive and robust strategy to limit aSyn protein burden, setting the stage for potential therapeutic interventions for a-Synucleinopathies.

Our original hypothesis was rooted in the understanding that endogenous aSyn plays a pivotal role in the seeding pathological processes that govern the propagation of a-Synucleinopathies. Therefore, we sought to assess the potential mitigating impact of aSyn downregulation on the pathological manifestations induced by the exogenous inoculation with human recombinant aSyn PFFs. Previous studies utilizing RNAi-based approaches, including shRNAs, siRNAs and microRNAs have shown that partial silencing of *Snca* gene reduces aSyn aggregation, preserves dopaminergic neurons and improves motor function in various experimental PD-like models [31]. However, initial studies achieving robust *Snca* down-regulation have raised concerns regarding the potential therapeutic utility of this approach due to neurotoxicity, inflammation and dopaminergic loss observed in animal studies [32]. Previous reports revealed that moderate aSyn silencing (e.g., 30–50% reduction) could evoke therapeutic benefits without adverse effects [33],[34]. More specifically, early reports demonstrated successful *Snca* silencing using lentiviral and AAV-based delivery of shRNA and ribozymes, which was accompanied by increased dopaminergic cell survival in both *in vitro* and *in vivo* models [35], [36]. In particular, a prior investigation conducted in the MPTP PD mouse model revealed an approximately 70% reduction in dopaminergic cell numbers, following MPTP treatment. This reduction was mitigated in animals injected stereotactically with sh-aSyn ribozymes-carrying rAAV vector, thereby rescuing a significant portion of the cell death and ameliorating subsequent motor impairment [36]. Furthermore, a reduction of the protein by approximately 35%, achieved through AAV2 intracerebral injection, exhibited notable neuroprotective effects [34]. Specifically, the AAV-sh-aSyn group demonstrated enhanced dopaminergic terminals in the striatum and reduced dopaminergic cell death induced by chronic rotenone exposure. The improved integrity of the nigrostriatal pathway was evident, as animals treated with sh-aSyn showed diminished motor deficits in behavioral tests. MicroRNAs like miR-7 and miR-153 also displayed neuroprotective effects in rodent a-Synucleinopathy models [37]. More recent advances, including AAV-mediated delivery of RNAi have refined this approach, demonstrating efficient *Snca* silencing and amelioration of aSyn-related pathology, while minimizing risks of neuroinflammation and transduction-related toxicity [38]. Furthermore, ASO-mediated aSyn down-regulation has been shown to attenuate the spread of pathological protein aggregates across neural networks, suggesting a potential neuroprotective role in mitigating PFF-induced neurodegeneration [39]. These studies underscore the potential of partial aSyn reduction as a safe, effective strategy for PD therapy.

In agreement with the latter studies, our data demonstrates the efficacy of partial *Snca* silencing in mitigating haSyn PFF-evoked pathology and relevant behavioral deficits, utilizing the non-invasive AAV-PHP.eB viral vectors. By achieving a balance between efficacy and safety, we demonstrate the efficient and sustained down-regulation of the endogenous mouse aSyn protein, utilizing two different AAV-PHP.eB constructs, harboring *Snca*-targeting shRNA or miRNA sequences. Both approaches curtailed aSyn protein levels, with the miRNA-targeting AAV-PHP.eB exhibiting improved knockdown efficacy (63% vs. 48%). This reduction was accompanied with mitigation of pSer129-aSyn-related pathology across multiple brain regions (including the substantia nigra), preservation of the nigral dopaminergic neurons and improved nigrostriatal dopaminergic terminal integrity and motor performance. Critically, these outcomes were achieved without evident off-target neurotoxicity or overall systemic adverse effects. Moreover, our results indicate a sustained expression of these AAVs accompanied by efficient down-regulation of endogenous aSyn protein levels and related amelioration of nigrostriatal dopaminergic system deficits at 5 months post-injection, thereby fortifying the utility of the employed viral vectors as enduring platforms for gene delivery. It is imperative to note, however, that the current findings on the 5-month cohort are contingent upon a limited number of animals (n=3/group), rendering them indicative and preliminary. Subsequent analyses with an expanded subject pool are requisite to substantiate and enhance the robustness of these outcomes.

AAV-mediated therapies have made a remarkable impact on treating neurological disorders, with AAV9 as a leading vector due to its ability to cross the BBB and reach CNS tissues [40]. However, achieving a broad and efficient spread throughout the CNS often requires high doses, which raises concerns about potential immune reactions and toxicity. To tackle this, researchers have developed AAV9 variants like AAV.PHP.B and AAV.PHP.eB, which demonstrate improved BBB penetration and CNS targeting in mice when administered peripherally. We selected to utilize AAV.PHP.eB viruses for their advantages over other viral vectors, including superior transduction efficiency compared to AAV.PHP.B and AAV9 [41]. The engineered PHP.eB capsid enables efficient transduction across diverse cell types, including neurons [40] rendering it ideal for the scope of our study. Additionally, its ability to cross the BBB allows intravenous administration, reducing infection risks and achieving widespread brain transduction, unlike localized injections [42]. Illustrative examples of the efficacious use of PHP.eB AAVs as gene delivery platforms are their application in the overexpression of the lysosomal enzyme β-glucocerebrosidase using AAV.PHP.B vectors in the A53T mouse model of PD [43] and in genetic models of Fragile X [44] and Angelman syndromes [34]. Such data highlight the adaptability and promise of these viral vectors as invaluable tools for advancing our comprehension of intricate biological mechanisms and laying the groundwork for innovative therapeutic interventions.

In summary, the findings from this study suggest that targeting the endogenous aSyn protein levels holds promise as a potential avenue for further exploration in the context of novel therapeutic strategies for PD and related synucleinopathies. This study achieved a significant milestone by non-invasively delivering shRNAs or microRNAs against endogenous murine *Snca* using the innovative AAV.PHP.eB, resulting in sustained and adequate expression of the transgene in an animal model that currently appears to possess the best face and construct validity among available PD models. The downregulation of endogenous aSyn demonstrated its efficacy in impeding the propagation of pathological aggregates into synaptically and anatomically connected brain regions. Additionally, it ameliorated the discernible neuronal demise observed in the nigrostriatal pathway.

Despite how appealing the reduction of aSyn protein levels currently is, the question of what constitutes a safe and effective degree of aSyn knockdown still remains unknown. Enhancing our understanding of the impact of aSyn downregulation on the neuronal system’s physiology is essential for maximizing its potential in combating a-Synucleinopathies [45]. Advancements in gene delivery vehicles, coupled with a deeper comprehension of the inherent mechanisms governing aSyn propagation, aggregation, and dissemination, may bring us closer to the ongoing quest for effective therapies for those afflicted by this condition.

### Limitations of the study

While phosphorylation of aSyn at residue Ser129 can serve as a robust marker for aSyn inclusion pathology, it should be interpreted cautiously. First, various cellular stresses or insults can significantly elevate the phosphorylation state of multiple proteins, including pSer129-aSyn without necessarily leading to inclusion formation [46]. Moreover, despite reports on the specificity and performance of the pSer129-aSyn antibody EP1536Y utilized in the current study for detecting aSyn aggregates in the brain tissue in the PFF model [47], some degree of cross-reactivity with other proteins and post-translational modifications in their vicinity cannot be ruled out. Thus, a secondary validation step is essential, which may involve the application of an alternative technique, such as proximity ligation assay (PLA), to reinforce the obtained results.

## Conclusions

This study highlights AAV-PHP.eB-mediated *Snca* silencing as a powerful strategy for mitigating aSyn pathology and functional impairments in a preclinical murine model of PD. By bridging mechanistic insights with therapeutic development, this study lays the groundwork for advancing aSyn-targeted interventions in PD and related synucleinopathies. Further optimization and validation in higher-order models will be critical for clinical translation. In addition, given the aforementioned promising characteristics of PHP.eB AAV, its potential use as a gene delivery tool for addressing aSyn-related dopaminergic neurodegeneration warrants closer examination. This becomes particularly pertinent considering emerging research indicating the significant correlation between aSyn protein load and dopaminergic neurotoxicity. Overexpression of aSyn caused by duplication or triplication of the gene, per se can lead to dopaminergic neurodegeneration by potentiating the formation of the oligomeric aggregates and eliciting the onset of PD. Even mild elevation of the protein levels is deleterious to neurons [48] Newer studies posit that there is a strong correlation between aSyn expression and regional vulnerability [11]. Curiously, despite the potential therapeutic value, the down-regulation of aSyn has not been extensively explored, and there is limited available data regarding the effectiveness of gene silencing and the vehicles employed for this purpose.

## Declarations

### Ethics approval and consent to participate

The animal study protocol was approved by the Ethical Committee for Use of Laboratory Animals of the Biomedical Research Foundation Academy of Athens (License No. 478434) and in accordance with the ARRIVE guidelines and the EU Directive 2010/63/EU for animal experiments.

### Consent for publication

Not applicable.

### Availability of data and material

All the raw data that support the findings of this study are available from the corresponding author, upon request.

### Authors’ contributions

Following the CRediT (Contributor Roles Taxonomy) author statement, the role(s) of each author are presented here: *Conceptualization:* Leonidas Stefanis, Maria Xilouri; *Data Curation:* Maria Fouka; *Formal Analysis:* Maria Fouka, Iraklis Tsakogias, Elena Giallinaki; *Funding acquisition:* Leonidas Stefanis, Maria Xilouri, George Tofaris; *Investigation:* Maria Fouka, Iraklis Tsakogias, Elena Giallinaki, Athanasios Stavropoulos; *Methodology:* Maria Fouka, Iraklis Tsakogias, Elena Giallinaki, Athanasios Stavropoulos, Maria Xilouri; *Project administration:* Leonidas Stefanis, Maria Xilouri; *Resources:* Christiane Volbracht, Louis De Muynck, Ronald Melki, Leonidas Stefanis, Maria Xilouri; *Software:* Elena Giallinaki, Iraklis Tsakogias; *Supervision:* Leonidas Stefanis, Maria Xilouri; *Validation:* Maria Fouka, Iraklis Tsakogias, Elena Giallinaki; *Visualization* Maria Fouka, Iraklis Tsakogias, Elena Giallinaki; *Writing – original draft:* Maria Xilouri, Maria Fouka, Iraklis Tsakogias; *Writing – review & editing:* Maria Xilouri, Leonidas Stefanis. All authors commented on previous versions of the manuscript. All authors read and approved the final manuscript.

## Acknowledgements

We would like to acknowledge the technical support of the Animal Housing facility personnel and Dr. Stamatis Pagakis, Anastasios Delis and Eleni Rigana of the Bioimaging Unit at BRFAA.

## Authors’ information (optional)

