## Supplementary Figures for "*In vivo* validation of novel non-invasive PHP.eB AAVs as a potential therapeutic approach for alpha-Synucleinopathies"

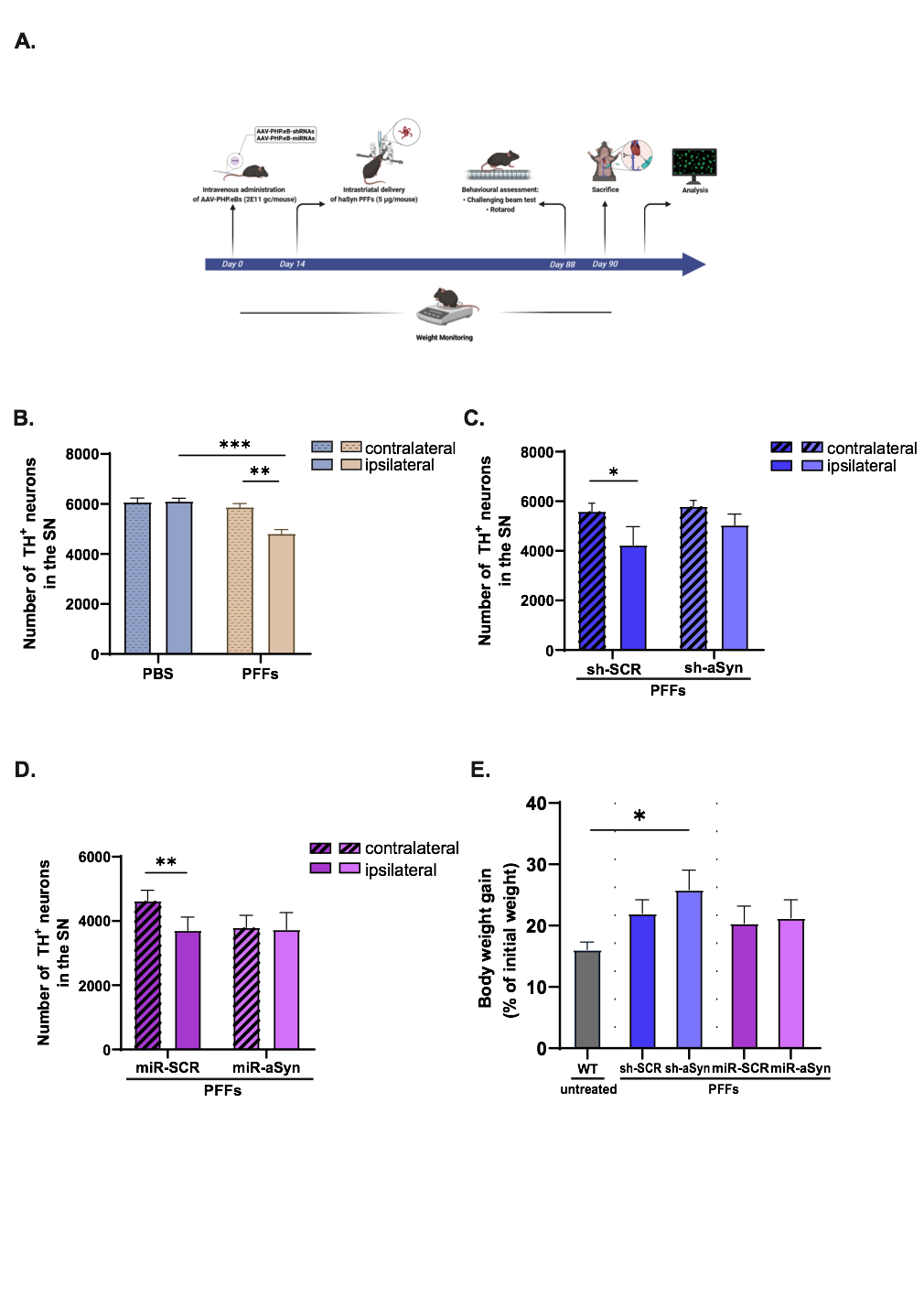

**Supplementary Figure 1. Systemic delivery of PHP.eBs does not affect dopaminergic neuron number and animal body weight. (A)** Schematic overview delineating the experimental design implemented. **(B-C)** Stereological counting of the dopaminergic nigral neurons reveal similar numbers at the contralateral sides of the PBS/PFF group (B) and sh-SCR or sh-aSyn AAV+PFFs-injected groups (C) further ratifying the non-toxic profile of the viruses utilized in this study. (**D**) Estimation of dopaminergic nigral neurons at the contralateral and ipsilateral sides of miR-SCR and miR-aSyn PHP.eB AAVs, using the Imaris 3 software, revealing similar TH^+^ neurons in both sides. Comparisons were made using two-way ANOVA, Šidák correction, asterisks indicate significance (*p < 0.05; ** p < 0.01; *** p < 0.001, n = 3-5 animals/group). (**E**) Weight gained in untreated (WT), shRNA- or miRNA- PHP.eB-PFF-treated animals illustrated as percentage of their initial body weight recorded at the first measurement (n = 5-7 for the WT and 8 animals /AAV-PHP.eB-treated group, Unpaired t-test).

**Supplementary Figure 2**. **Stable GFP-PHP.eB AAV transgene expression within the nigrostriatal pathway at 5 months post-systemic administration**. **(A)** Representative immunofluorescence midbrain images with antibodies against TH (red) and GFP (green) depicting the expression of GFP-tagged AAV.PHP.eB sh-SCR within mouse TH^+^ dopaminergic nigral neurons, 5 months post i.v. injection. DAPI was used as nuclear marker. Scale bar: 100 μm. **(B)** Quantification of the GFP^+^/TH^+^ transduced dopaminergic neurons 5 months post i.v. injection. **(C-F)** Representative pSer129-aSyn^+^ DAB immunostainings and respective quantifications of the percentage of pSer129-aSyn^+^ area in the striatum (C-D) and substantia nigra (E-F), in all groups using antibody against pSer129-aSyn. Scale bar: 100 μm. (**G-H)** Representative TH DAB immunostaining striatal (C-D) and midbrain (E-F) sections and quantification (G) and quantification of the TH^+^ fiber density relative to the corresponding contralateral hemisphere. Improved TH^+^ density in the ipsilateral hemisphere of the sh-aSyn+PFFs animals as compared to the sh-SCR+PFFs treated ones (** p < 0.01). Statistical significant reduction of TH density in ipsilateral hemispheres of sh-SCR+PFFs. Comparisons were made using two-way ANOVA, Šidák correction, asterisks indicate significance (*** p < 0.001; **** p < 0.0001; * p < 0.05 animals/group). **(I)** Evident motor deficits in AAV-PHP.eB sh-SCR+PFFs injected animals (*p < 0.05, n = 3 animals/group, One-way ANOVA followed by Tukey’s post hoc test) assessed at 5 months post-AAV administration. Quantifications of errors per step measured by the challenging beam test are shown.
